# A FOXN1 mutation competitively displaces wild-type FOXN1 from higher-order nuclear condensates to cause immunodeficiency

**DOI:** 10.1101/2021.03.28.437237

**Authors:** Ioanna A. Rota, Adam E. Handel, Fabian Klein, Stefano Maio, Fatima Dhalla, Mary E. Deadman, Stanley Cheuk, Joseph A Newman, Yale S. Michaels, Saulius Zuklys, Nicolas Prevot, Philip Hublitz, Philip D. Charles, Athina Soragia Gkazi, Eleni Adamopoulou, Waseem Qasim, E. Graham Davies, Celine Hanson, Alistair T. Pagnamenta, Carme Camps, Helene M. Dreau, Andrea White, Kieran James, Roman Fischer, Opher Gileadi, Jenny C. Taylor, Tudor Fulga, B. Christoffer Lagerholm, Graham Anderson, Erdinc Sezgin, Georg A. Holländer

**Affiliations:** Department of Paediatrics and the MRC Weatherall Institute of Molecular Medicine, University of Oxford, Oxford, UK; MRC Functional Genomics Unit, Department of Physiology, Anatomy and Genetics, University of Oxford, Oxford, UK; Structural Genomics Consortium, University of Oxford, ORCRB, Roosevelt Drive, Oxford, UK; MRC Weatherall Institute of Molecular Medicine, Genome engineering services, Radcliffe Department of Medicine, University of Oxford, Oxford, UK; Genome Engineering and Synthetic Biology Unit, MRC Weatherall Institute of Molecular Medicine, University of Oxford, Oxford, UK; Target Discovery Institute, University of Oxford, Oxford, OX3 7FZ, UK; Great Ormond Street Hospital & Great Ormond Street Institute of Child Health, University College London; Department of Pediatrics, Baylor College of Medicine, Houston, TX, USA; National Institute for Health Research Biomedical Research Centre, Oxford, UK; Wellcome Centre for Human Genetics, University of Oxford, Oxford, OX3 7BN, UK; Department of Oncology, University of Oxford, Oxford, OX3 7DQ, UK; Institute for Immunology and Immunotherapy, Medical School, University of Birmingham, B15 2TT, UK; Wolfson Imaging Centre Oxford, MRC Weatherall Institute of Molecular Medicine, University of Oxford, Headley Way, Oxford OX3 9DS, UK; MRC Human Immunology Unit, MRC Weatherall Institute of Molecular Medicine, University of Oxford, Oxford, UK; Science for Life Laboratory, Department of Women’s and Children’s Health, Karolinska Institutet, Solna, Sweden; Paediatric Immunology, Department of Biomedicine, University of Basel and University Children’s Hospital Basel, Basel, Switzerland; Department of Biosystems Science and Engineering, ETH Zurich, Basel, Switzerland

**Keywords:** Forkhead box N1 transcription factor, immunodeficiency, thymus

## Abstract

The transcription factor FOXN1 is a master regulator of thymic epithelial cell development and function. Here we demonstrate that FOXN1 expression is differentially regulated during organogenesis and participates in multi-molecular nuclear condensates essential for the factor’s transcriptional activity. FOXN1’s C-terminal sequence regulates the diffusion velocity within these aggregates and modulates the binding to proximal gene regulatory regions. These dynamics are significantly altered in a patient with a mutant FOXN1 which is modified in its C-terminal sequence. This mutant is transcriptionally inactive and acts as a dominant negative factor displacing wild-type FOXN1 from condensates and causing athymia and severe lymphopenia in heterozygotes. Expression of the mutated mouse ortholog, selectively impairs mouse thymic epithelial cell (TEC) differentiation revealing a gene dose dependency for individual TEC subtypes. We have therefore identified the cause for a primary immunodeficiency disease and determined the mechanism by which this FOXN1 gain-of-function mutant mediates its dominant negative effect.

---

The thymus microenvironment promotes the development of naïve T cells with a repertoire purged of “self” specificities and poised to react to potentially injurious “non-self” threats. Thymic epithelial cells (TEC) constitute the major component of the thymic stroma and can be categorised into separate lineages and states based on their specific molecular, structural, and functional characteristics[1]. TEC differentiation, maintenance, and function critically rely on the transcription factor FOXN1 [2–8]. FOXN1 is a member of the forkhead box (FOX) family of transcription factors and recognises a 5 base-pair consensus sequence (GACGC) via its centrally located DNA-binding domain (the Forkhead domain, FKH)[2]. Direct and water-mediated contacts via residues in an α-helix inserted in the DNA major groove dictate the binding of the FKH to its consensus binding site [9]. In addition, the first 154 N-terminal amino acids for FOXN1 are required for normal TEC differentiation and the acidic activation domain in the C-terminal region is required for target gene transcription [10, 11].

Many transcription factors operate within large nuclear biomolecular condensates, which resemble aggregates formed by liquid-liquid phase separation. The lack of a membrane surrounding nuclear condensates allows rapid exchange of components with the nucleoplasm [12]. Nuclear condensates can impact chromatin architecture by maintaining the heterochromatin domain[13] or have been shown to drive gene activation through the assembly of transcriptional complexes at enhancer-rich gene regulatory regions[14]. Proteins in membrane-less nuclear organelles characteristically include intrinsically disordered regions (IDRs) [14–17]. FOXN1 has several IDRs, although whether or how it might function within a nuclear biomolecular condensate under physiological and pathological conditions has remained largely undefined.

Autosomal recessive mutations of *FOXN1* that result either in a premature stop in translation (p.R255X) [5, 18], a loss of DNA binding (p.R320W) [18], or a frameshift and premature truncation (p.S188Afs*114) [19] give rise to a rare, phenotypically unvarying form of congenital severe combined immunodeficiency known as lymphoid cystic thymic dysgenesis (a.k.a. “nude” phenotype; ORPHA169095). In addition to the absence of thymic tissue and the consequent lack of T cells, the syndrome is further characterized by alopecia universalis and nail dystrophy as a result of a lack of functional FOXN1 expression in the ectoderm [4]. In contrast to the complete constitutive absence of FOXN1 expression, reduced expression of FOXN1 in fetal and young mice causes only a transient hypoplasia of the thymus, [20] whereas lower FOXN1 concentrations in older animals are associated with premature thymic involution [8].

Together with recent clinical observations of individuals with heterozygous FOXN1 mutations[21], these findings suggest that the formation and maintenance of the thymus appears to be sensitive to small changes in FOXN1 availability, the consequences of which can be severe for the immune system. However, the molecular mechanism by which FOXN1 exerts its precise transcriptional function, and whether this might involve nuclear biomolecular condensates has remained unclear [22, 23].

## Results

### Athymia and T cell lymphopenia caused by heterozygous FOXN1 mutants

We have identified a FOXN1 variant with a single base-pair deletion in exon 7, NM_001369369 :c.1370delA (p.H457Pfs*93), in three individuals of a single kindred. This variant caused a frame shift resulting in a “scrambled” sequence of 92 amino acids (aas) and a premature stop codon at aa 549 of FOXN1, and is designated thereafter as Δ550 FOXN1 (Figure 1a and Supplementary Figure 1a). The affected individuals were heterozygous for the Δ550 FOXN1 mutation but presented clinically with athymia and T cell lymphopenia, the latter characterized in the index case by the absence of T cell receptor (TCR) excision circles (TRECs, i.e. small circles of DNA created by rearrangement of TCR genes used as a surrogate marker to assess recent thymic T cell output) and a significantly reduced TCR repertoire diversity (Supplementary Figure 1b and 1c).

**Figure 1:**
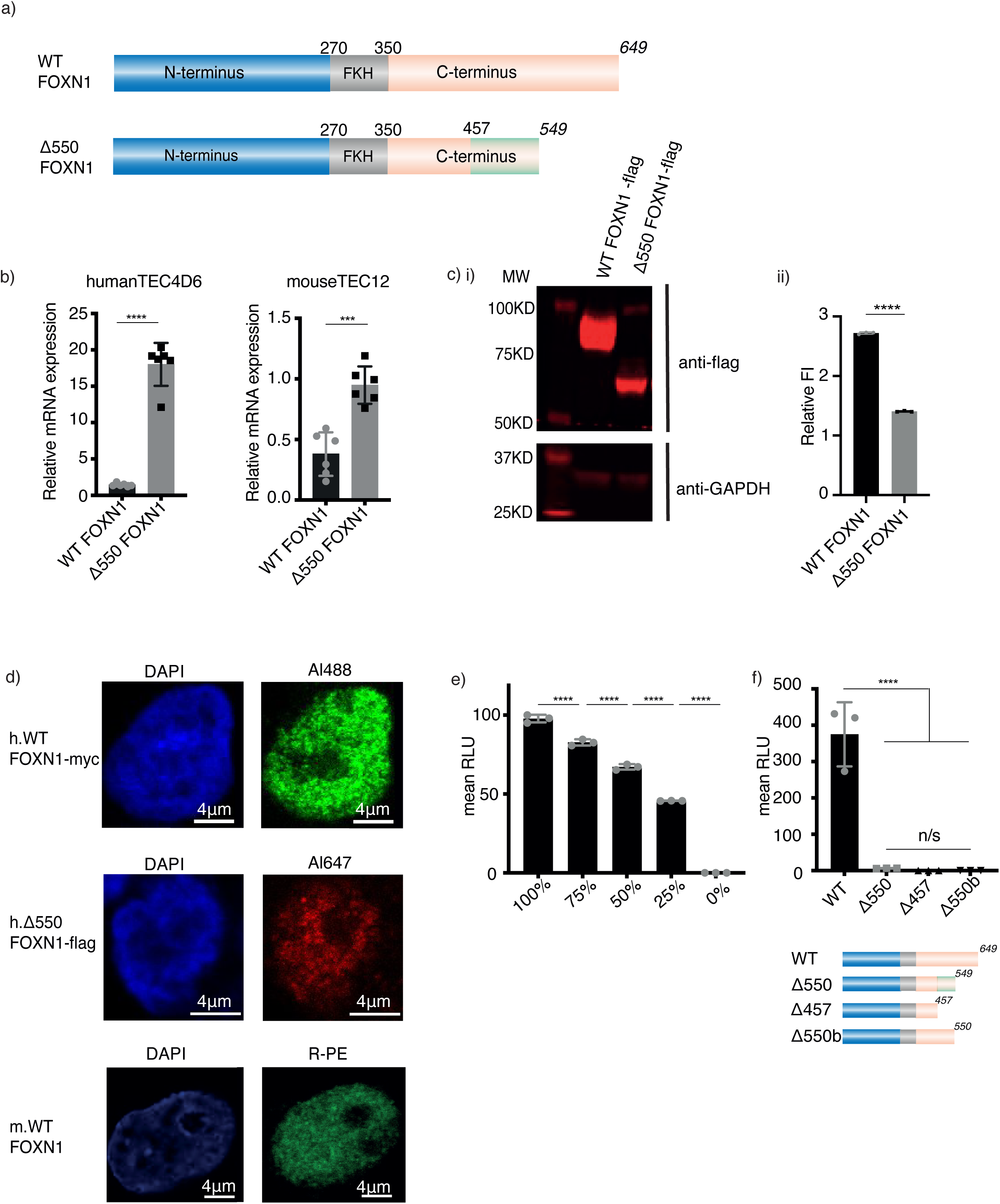
The Δ550 FOXN1 mutation lacks transcriptional activity. (a) Schematic representation of wild-type (WT) FOXN1 and the Δ550 FOXN1 mutant. FKH: forkhead domain. Numbers represent amino acids, numbers in italics indicate the position of the stop codon and the green box indicates the scrambled amino acid sequence. (b) Relative wild-type and Δ550 FOXN1 mRNA expression compared to GAPDH in transfected human 4D6 and mouse TEC1.2 cells. (c) Western blot (i) and quantification (ii) of FLAG-tagged wild-type and Δ550 FOXN1 protein in 4D6 cells. Fluorescence intensity (FI) was compared to that of GAPDH after background subtraction. (d) Confocal microscopy of the nuclear localization of untagged wild-type (WT, bottom panel), myc-tagged WT (middle panel) and FLAG-tagged Δ550 FOXN1 in 4D6 cells (top panel) using indirect and direct immunofluorescence, respectively, and DAPI counter staining.. (e) Linear relationship between wild-type FOXN1 gene dose and transcriptional activity in transfected 4D6 cells expressing a luciferase reporter construct. Luciferase activity was quantified as arbitrary relative light units (RLU). (f) Top: Transcriptional activity of wild-type and mutant FOXN1 in 4D6 cells as measured by luciferase reporter assay. Bottom: Schematic representation of FOXN1 variants with scrambled amino acid sequences shown as a light green box. The data is representative of 1 (panel d), 2 (panel c), 3 (panels e, f) and 6 (panel b) independent experiments with three technical replicates each. Each symbol represents data from an individual experiment. Mean value and SD are shown and statistically compared by two-tailed unpaired t-test (panel b, c, f) and ANOVA (e); p-values: ≥ 0.05 (ns), 0.0002 (***), <0.0001(****). See also Figure S1.

To characterize the Δ550 FOXN1 variant, we engineered a single-base pair deletion at position 1370 in the human wild-type *FOXN1* sequence and added either a 3’ flag or myc tag to both the wild-type and mutant *FOXN1* genes. Wild-type and Δ550 *FOXN1* sequences were expressed using the same vector in human (TEC4D6) and mouse (TEC1.2) TEC lines. Higher mRNA levels were observed for the mutant compared to wild-type (Figure 1b) indicating that the Δ550 FOXN1 mRNA escaped non-sense mediated decay. In contrast, transfected TEC4D6 cells contained lower Δ550 FOXN1 protein levels when compared to the wild-type protein, suggesting that translation efficiency and/or stability were reduced for the mutant protein (Figure 1c).

To determine Δ550 FOXN1’s subcellular localization, TEC4D6 cells were transfected with either wild-type or mutant FOXN1 and the location of each variant was determined by immunostaining. Both wild-type and Δ550 FOXN1 were present in the nucleplasm, expressed in a speckled pattern, and were excluded from the nucleolus, as was seen for untagged FOXN1 detected using an anti-FOXN1 antibody (Figure 1d). The FOXN1 nuclear expression pattern is similar to transcription factors that are part of phase-separated multimolecular condensates [14].

The transcriptional activity of wild-type and Δ550 FOXN1 were assayed in TEC4D6 cells co-transfected with a reporter plasmid containing a luciferase gene under the transcriptional control of the FOXN1-specific *Psmb11* promoter, which normally controls β5t expression in TEC [2]. Wild-type FOXN1 activated the luciferase gene in a gene dosage sensitive manner (Figure 1e) whereas Δ550 FOXN1 failed to activate luciferase gene expression (Figure 1f). Luciferase activity was also not observed in transfectants of C-terminally truncated wild-type FOXN1 variants, where a stop codon was introduced either at amino acid (aa) 457 (the position of the Δ550 FOXN1 missense mutation) or 550 (the Δ550 FOXN1 premature stop codon) (Figure 1f), consistent with the C-terminal domain playing a critical role in FOXN1-mediated gene activation [10, 11]. Although Δ550 FOXN1 apparently behaved as a loss-of-function mutant for transcriptional activation, its heterozygosity causing athymia was not explained by these results.

### *Δ505 Foxn1* heterozygous mice show TEC defects

To analyze the impact of Δ550 FOXN1 heterozygosity on thymus development and function, mice were generated that have a single nucleotide deletion, orthologous to that observed in the index patient, at position 1370 of the murine *Foxn1* gene. The ensuing frame shift resulted in a scrambled protein sequence starting at aa 457 and a premature stop codon at aa 505 (named Δ505; Supplementary Figure 2a). Five- and 16-week old male mice (but not embryos) heterozygous for the Δ505 FOXN1 mutation (FOXN1^WT/Δ505^) had a thymus with significantly reduced overall cellularity when compared to age-matched wild-type littermates (FOXN1^WT/WT^) (Figure 2a, Supplementary Figure 2). While their total TEC counts were comparable with wild-type littermates (Figure 2b), the postnatal FOXN1^WT/Δ505^ mice displayed changes in the relative number of flow-cytometrically defined TEC subpopulations (Figure 2 c and 2d). The relative frequency of TEC (EpCAM+CD45-) and cortical (c) TEC (EpCAM+Ly51+UEA-) was increased in FOXN1^WT/Δ505^ mice at both time points (Figure 2b and 2c, see also and Supplementary Figure 2b). In addition to a lower frequency of total medullary (m) TEC (EpCAM+Ly51-UEA+) at 16 weeks, the differentiation within that lineage revealed a partial maturational block. Fewer mature mTEC were detected after birth that either expressed the autoimmune regulator (AIRE) required for promiscuous gene expression (i.e. CD80^pos/hi^) or that had progressed beyond that specific developmental stage (a.k.a. post-AIRE, e.g. Tspan8+; Figure 2di-iii; see also Supplementary Figures 2c-e). Moreover, mTEC, and in particular cells with an immature phenotype expressed lower levels of MHC class II on their cell surface (Figure 2div and Supplementary Figure 2f). However, the frequency of TEC expressing full-length FOXN1 and the geometric mean fluorescence intensity of wild-type FOXN1 in TEC were reduced in FOXN1^WT/Δ505^ mice when quantified using an antibody that binds the C-terminus between amino acids 475 and 542 (Figure 2g) [24]. Both TEC frequency and cellularity were normal in age-matched FOXN1^WT/nu^ mice, i.e. mice heterozygous for the nude locus which encodes a FOXN1 mutation unable to bind to DNA (Supplementary Figure 2g), suggesting that the differences observed were not secondary to a reduction in *Foxn1* gene dosage.

**Figure 2:**
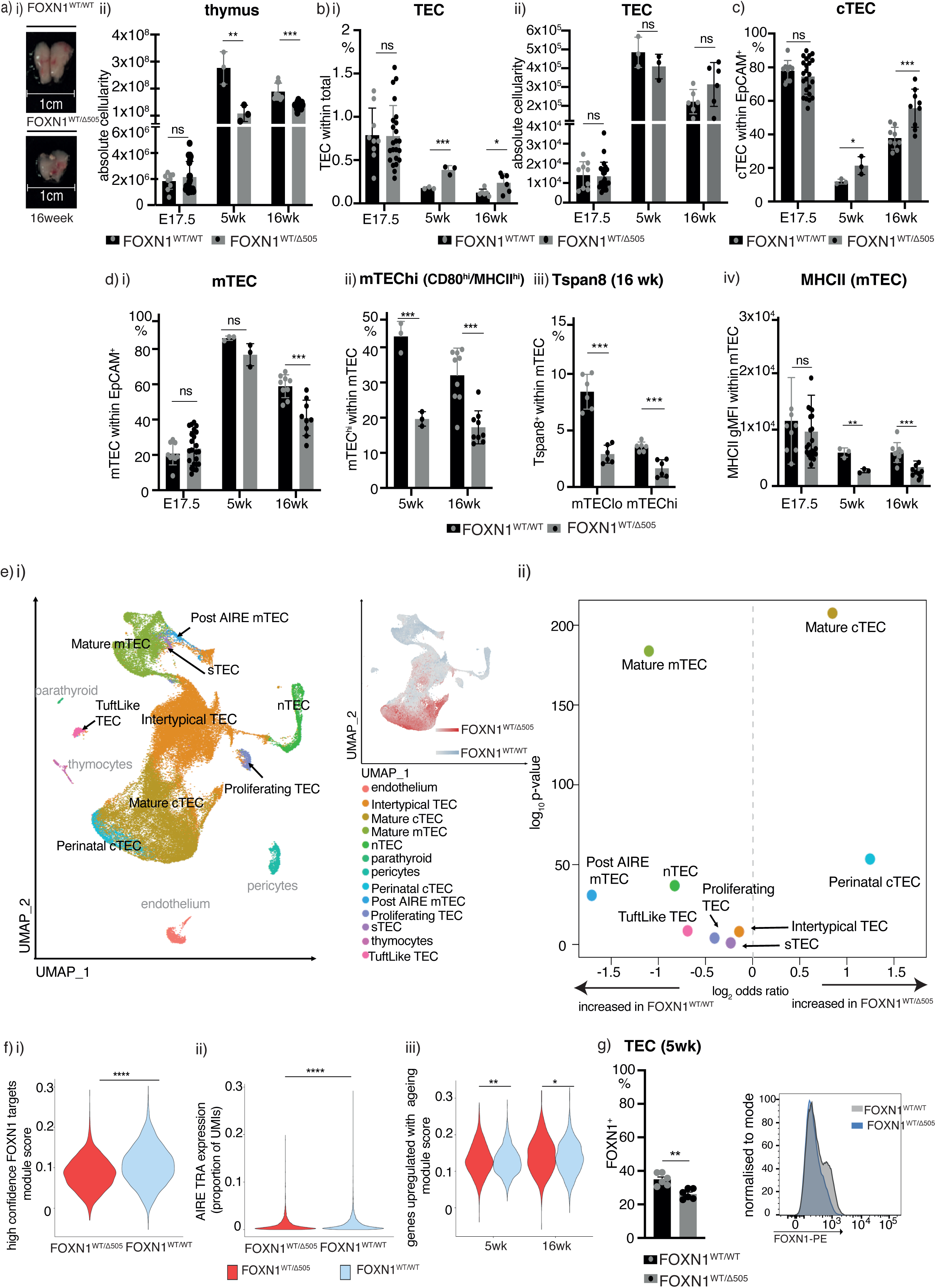
Heterozygous Δ505 FOXN1 expression in mice impairs TEC differentiation and promiscuous gene expression. Comparison of gender-matched FOXN1^WT/WT^ and FOXN1^WT/Δ505^ mice at indicated ages: (a) Thymus gross anatomy and (ii) absolute cellularity. (b) (i) Relative and (ii) absolute TEC cellularity. (c) Relative cTEC cellularity. (d) Relative cellularity of (i) total mTEC and (ii-iii) their phenotypically distinct subpopulations; (iv) level of MHCII expression on the surface of total mTEC. (e) Single cell TEC analysis of FOXN1^WT/WT^ and FOXN1^WT/Δ505^ mice. (i) Uniform Manifold Approximation and Projection (UMAP) analysis of combined single TEC transcriptomic data with an inset showing local enrichment of TEC single cells by genotype (using the 200 nearest neighbours). (ii) Comparative analysis of TEC subtype frequencies. (f) Violin plots of data from mature mTEC: (i) Expression of FOXN1 high confidence target genes, (ii) the proportion of unique molecular identifiers (UMIs) assigned to AIRE-induced genes and (iii) the expression of genes upregulated in a tissue-independent manner throughout ageing (g) Comparison of (i) frequency of FOXN1^+^ cells and (ii) FOXN1 expression levels in TEC from gender-matched FOXN1^WT/WT^ and FOXN1^WT/Δ505^ mice at 5weeks. Each symbol in panels a-d,g represents data from an individual wild-type or mutant mouse. The flow cytometric gating strategies for panels a-d are shown in Supplementary Figure 3 and for panel g in Supplementary Figure 8. The data in panels a-d are from 4 (E17.5), 1 (week 5) and 3 (week 16) independent experiments, for panel g from 2 independent experiments with each at least 3 wild-type and mutant male mice. Mean value and SD are shown in bar graphs and were calculated two-tailed unpaired t-test; 0.05 (ns), 0.0332(*), 0.0021(**), 0.0002 (***). Data in panel e is from 18,637 and 36,943 cells isolated from 3 FOXN1^WT/WT^ and 3 FOXN1^WT/Δ505^ mice each at 5 and 16 weeks of age, respectively. Data in panel e ii displays log_2_ odds ratio and -log_10_ p-values using Fisher’s exact test. The data shown in panel f are from 3 FOXN1^WT/WT^ and 3 FOXN1^WT/Δ505^ mice: 6,687 mature mTEC, p<0.05(*), p <0.01(**), p<0.0001(****). See also Figure S2, S3 and table S1 and S2

To further probe the consequences of Δ505 *Foxn1* heterozygosity for TEC differentiation, we assessed the transcriptome of single epithelial cells using a recently published data set as reference to infer distinct TEC subtypes [1]. The composition of the TEC scaffold was significantly changed in mutant mice, affecting the frequency of all but one of the TEC subtypes (i.e. sTEC; Figure 2e) with concomitant alterations in biological pathways relating to antigen presentation, cytokine responses, and cell proliferation (Supplementary table 1 and 2). These changes in TEC lineage maturation included an expansion both of perinatal cTEC, a subtype substantially reduced in wild-type mice 4 weeks of age and older, and mature cTEC. In contrast, the frequencies of almost all of the other TEC subtypes were reduced when compared to wild-type mice, with a particularly large reduction in mature and post-AIRE mTEC. The relative diversion of TEC into the cortical rather than the medullary lineage in FOXN1^WT/Δ505^ mice was substantial in intertypical TEC, as we observed significantly lower expression of *Krt5*, a marker of mTEC fate, and higher expression of *Prss16*, a marker of cTEC fate (all p < 0.0001) (Supplementary Figure 2h). This alteration in the mTEC compartment impaired both the expression of FOXN1 target genes (Figure 2fi) [2]and AIRE-controlled tissue-restricted antigens (Figure 2fii)[25]. Moreover, mature mTEC isolated from FOXN1^WT/Δ505^ mice at either 5- or 16-weeks of age showed evidence of accelerated ageing in comparison to controls [1, 26] thus suggesting that cells passing through the mTEC developmental block were under increased cellular stress (Figure 2fiii). In particular, mature mTEC from FOXN1^WT/Δ505^ mice showed increased expression of a gene associated with lysosomal injury (*Serpinba6*) but lower expression of a constituent of the immunoproteasome associated with alleviation of proteotoxic stress in mTEC (*Psmb10*) [27, 28].

We evaluated whether these epithelial changes affected thymopoiesis in FOXN1^WT/Δ505^ mice. At 5- and 16-weeks of age, thymocyte differentiation was generally comparable to that of wild-type mice (Figure 3a and Supplementary Figure 4). However, thymocyte negative selection was compromised in young and older FOXN1^WT/Δ505^ mice as fewer CD4+CD8+ (double positive, DP) and mature stage 2 single CD4-positive cells (M2: CD69^-^ MHC I^+^) and single CD8-positive cells at the stages M1 (CD69^+^MHC I^+^) and M2 underwent clonal deletion (Figure 3b). In addition, the frequency of NKT cells (Figure 3c) but not that of T_reg_ and γδ Τ cells was reduced in FOXN1^WT/Δ505^ mice (Supplementary Figure 4e,f). Thus, heterozygosity for a mouse orthologue of human D550 FOXN1 substantially impaired TEC differentiation and function.

**Figure 3.**
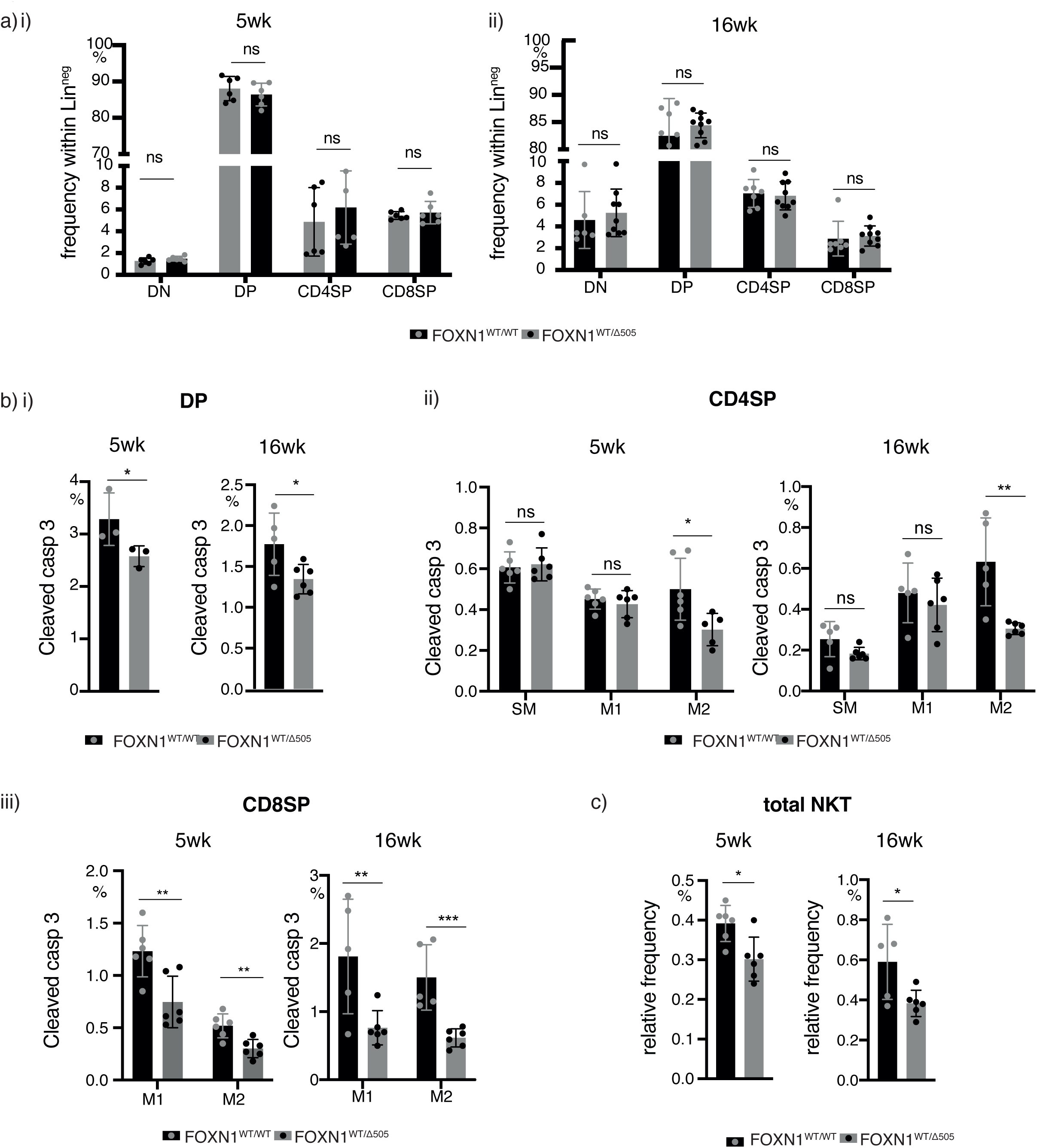
Heterozygous Δ505 FOXN1 expression impairs T cell clonal deletion at the population level. (a) Thymocyte differentiation in (i) 5 week-old and (ii) 16 week-old FOXN1^WT/WT^ and FOXN1^WT/Δ505^ mice. Live thymocytes negative for CD11b, CD11c, Gr1, CD19, CD49b, F4/80, NK1.1, TCRγδ, and Ter119 expression (designated Lin-) were stained for the surface expression of CD4 and CD8. Gating strategy in Supplementary Figure 3. (b) Age-specific analysis of the frequency of negatively selected (i) double positive (DP), (ii) CD4 single positive (CD4SP) thymocytes and (iii) CD8 single positive (CD8SP) thymocytes as assessed by the detection of cleaved caspase 3. The CD4SP cells were differentiated into semi-mature (SM: CD69^+^MHC I^low^), mature 1 (M1: CD69^+^MHC I^+))^) and mature 2 (M2: CD69^-^ MHC I^+^) thymocytes [51]. The CD8SP cells are distinguished into mature 1 (M1: CD69^+^MHC I^(+)^) and mature 2 (M2: CD69^-^ MHC I^+^) thymocytes. (c) Frequencies of NKT cells (TCRβ^+^, CD1d tetramer^+^) among thymocytes and [52]. Each symbol represents data from an individual wild-type or mutant mouse. Data is from 3 (panel a) and 2 (panels b,c) independent experiments, respectively. Mean value and SD are shown and statistically compared by two-tailed unpaired t-test. 0.05 (ns), 0.0332(*), 0.0021(**). See also Figure S3 and S4

### Δ550 FOXN1 is a dominant negative mutant

Haploinsufficiency caused by hypomorphic *FOXN1* variants have been associated with T cell lymphopenia and thymic hypoplasia/aplasia [21] and thus may also explain the clinical phenotype of individuals heterozygous for the transcriptionally defective Δ550 FOXN1 variant (Figure 1f). Alternatively, Δ550 FOXN1 could act as a dominant negative mutant interfering with the transcriptional activity of wild-type FOXN1. To test this hypothesis, we co-expressed wild-type and mutant Δ550 FOXN1 together with a FOXN1-specific luciferase reporter in TEC 4D6 cells [2]. Contrary to the gene-dose-dependent signal detected with wild-type FOXN1, co-expression of wild-type and variant FOXN1 substantially reduced luciferase activity (Figure 4a), identifying Δ550 FOXN1 as a dominant negative mutant. Similar results were obtained in a murine TEC cell line co-expressing mouse wild-type FOXN1 and the Δ505 FOXN1 mutant (Supplementary Figure 7a). Thus, a single nucleotide deletion generating the human Δ550 and mouse Δ505 mutations creates dominant negative variants of FOXN1.

**Figure 4:**
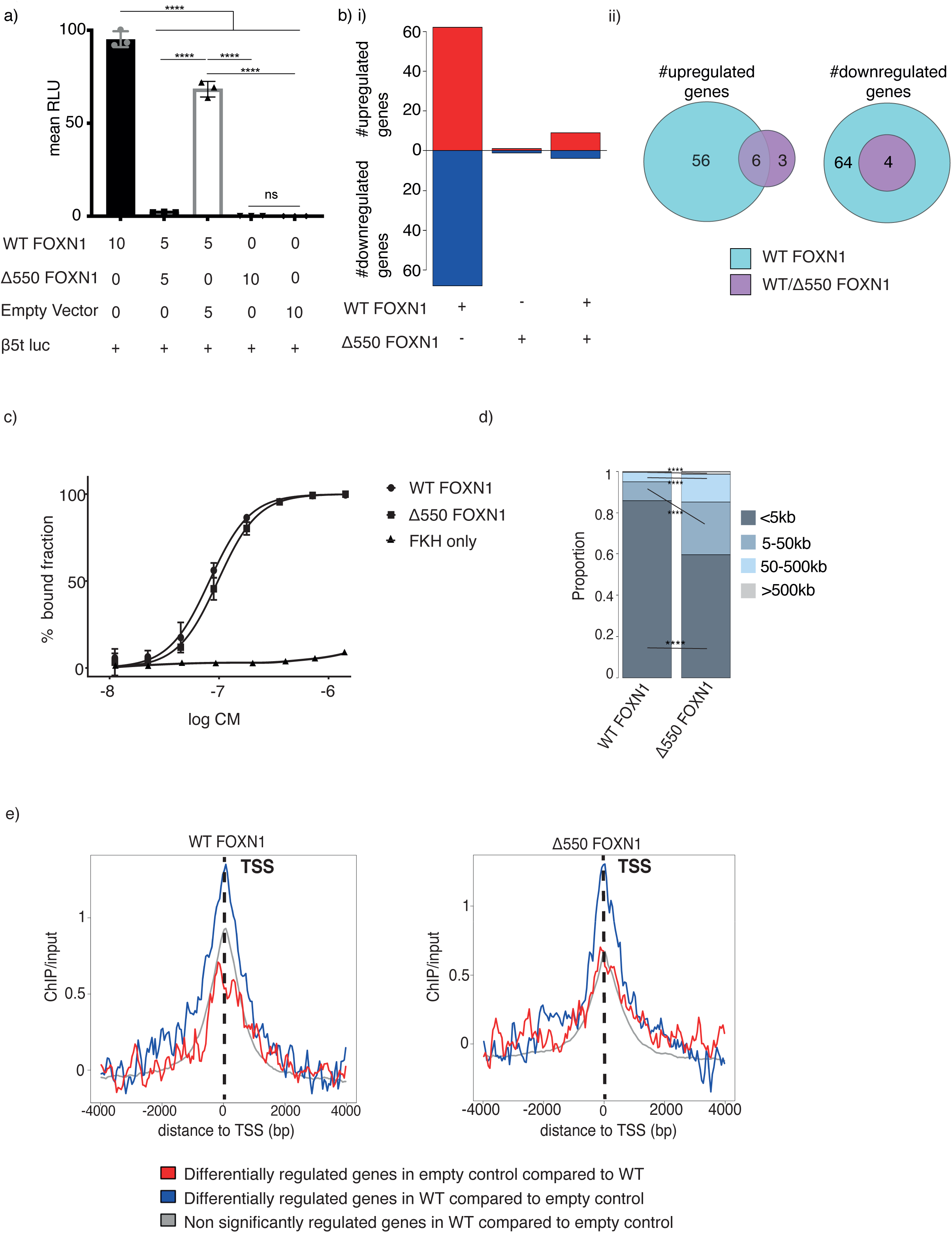
Δ550 FOXN1 is transcriptionally inactive and acts a dominant negative variant to compete with wild-type FOXN1 for DNA binding. (a) Expression of a luciferase reporter in 4D6 cells transfected with indicated expression vectors. Constitutive renilla expression was used in each transfectant as an internal control and reporter activity was measured as arbitrary relative light units (RLU) following correction. WT: wild-type. (b) Gene expression changes in 4D6 cells transfected to express wildtype and Δ550 FOXN1 either alone or in combination; (i) number of genes differentially expressed in comparison to control transfected cells, and (ii) changes in the number of genes commonly or separately changed in the presence of wild-type and Δ550 FOXN1. (c) Gel shift assay to determine DNA binding affinity of wild-type FOXN1, Δ550 FOXN1 and isolated FOXN1 forkhead domain (FKH) to a DNA sequence containing the FOXN1 binding motif. Quantification of bound fraction fitted to a standard binding isotherm. (d) Proportion and comparison of distances to the closest TSS for wild-type and Δ550 FOXN1 ChIP-seq peaks. (e) Wild-type (left) and Δ550 FOXN1 ChIP-seq peaks (right) flanking the transcriptional start site (TSS) of genes differentially regulated by wild-type FOXN1. The data shown is representative of 4 (a) and 3 independent experiments (c) each with at least 3 technical replicates. Panels b, d, and e display an independent experiment with 3 biological replicates each. Mean value and SD are shown and were and statistically compared by two-tailed unpaired t-test (panel a) and Fisher’s test (d): p-values: ≥0.05 (ns), <0.0001(****). See also Figure S5

To verify the dominant negative nature of Δ550 FOXN1, we analyzed gene expression profiles of the human TEC line 4D6 transfected with either wild-type or Δ550 FOXN1 alone, or a combination of the two (Figure 4b), using RNA-Seq. Wild-type FOXN1 up-regulated 62 genes and down-regulated 68 genes. In contrast, Δ550 FOXN1 did not up-regulate any genes and down-regulated only a single gene, confirming its lack of transcriptional activity. Co-expression of wild-type and Δ550 FOXN1 resulted in the up- or down-regulation of only a fraction of genes controlled by wild-type FOXN1 (6/62 [9.6%] and 4/68 [5.8%], respectively) (Figure 4b). The binding affinities of wild-type and Δ550 FOXN1 were statistically indistinguishable for a DNA probe containing two copies of the canonical GACGC motif (Figure 4c)[9]. ChIP-seq analysis of 4D6 cells expressing either wild-type FOXN1-FLAG or Δ550 FOXN1-FLAG identified 10,273 and 16,426 peaks, respectively (at an irreproducible discovery rate (IDR) of <0.05) with significant enrichment in their overlap (122.1-fold, p < 0.0001; Supplementary Figure 5a). FOXN1 binding occurred within a 5 kb window of the gene transcriptional start sites for 85.9% of sites bound by wild-type FOXN1and 59.7% occupied by Δ550 FOXN1 (Figure 4d). The consensus FOXN1-binding motif, GACGC (Supplementary Figure 5b), was identified by the motif analysis MEME-Chip [29] within wild-type FOXN1 ChIP-seq peaks as the top ranked candidate sequence whereas the same motif was identified at a lower frequency in Δ550 FOXN1 ChIP-seq peaks (63.1% *vs.* 49.2%, p < 0.0001). Hence, the majority of wild-type but significantly fewer mutant FOXN1 molecules bound to proximal gene regulatory regions recognizing the same DNA motif (p < 0.0001), indicating the C-terminal region of FOXN1 modulates both transactivation and DNA binding.

The FOXN1 ChIP-seq data were integrated with the RNA-Seq data, revealing that only genes controlled by wild-type FOXN1 were also enriched for FOXN1 ChIP-seq signals in 4D6 cells (Figure 4e). However, neither wild-type nor Δ550 peaks were enriched near the few genes differentially regulated by Δ550 FOXN1 (Supplementary Figure 5c). Taken together, these results support a mechanism by which Δ550 FOXN1 successfully competes with wild-type FOXN1 for DNA binding but due to its lack of a transactivation domain does not initiate transcription.

### Δ550 FOXN1 impairs condensate dynamics

Transcription factors control gene expression as either monomers, homodimers, heterodimers or oligomers [30]. To investigate whether FOXN1 operates either as a dimer or a multimer, 4D6 cells were co-transfected with two forms of wild-type FOXN1, each labeled with either a FLAG- or myc-tag. Immunopreciptation using an antibody directed against one tag also recovered FOXN1 labeled with the other tag, demonstrating the formation of FOXN1 homo-dimers or multimers. This complex formation was inhibited in the presence of the Δ550 variant (Figure 5a), thus demonstrating the physical impact of the mutant on FOXN1 multimer formation.

**Figure 5:**
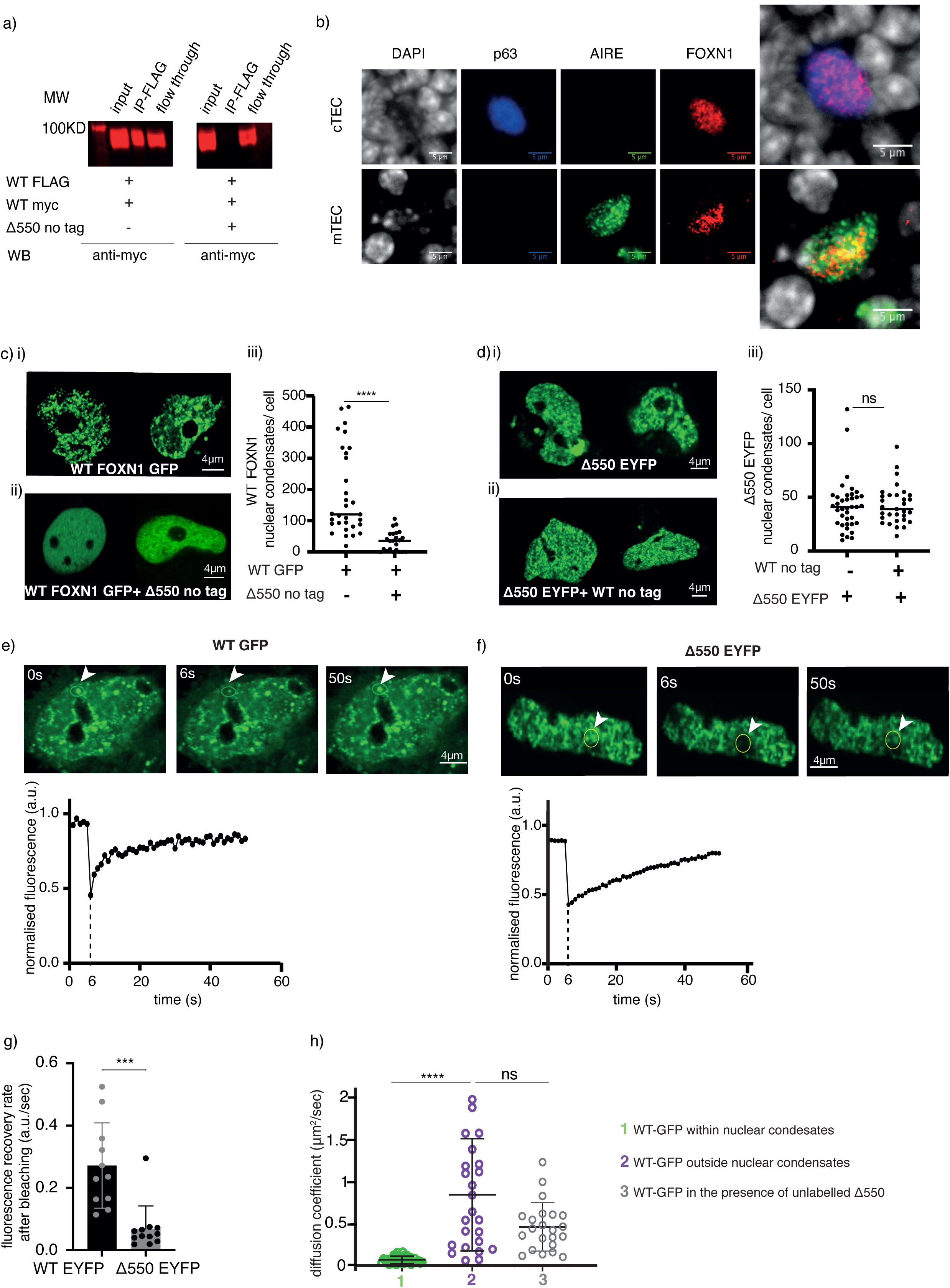
Δ550 FOXN1 disrupts the formation of wild-type FOXN1 multimers. (a) Anti-Flag immuno-precipitates and Western blot analysis of FLAG- and myc-tagged wild-type (WT) FOXN1 expressed in the absence or presence of Δ550 FOXN1 in 4D6 cells. Protein detection using anti-myc antibody. (b) Widefield microscopy showing FOXN1 nuclear condensates in primary mouse TEC. FOXN1 (red), AIRE (green) and p63 (blue) direct immunofluorescence and DAPI counterstain of thymic tissue sections from 1 week old wild-type mouse. AIRE-positive mTEC do characteristically not express p63. (c) Confocal microscopy showing nuclear condensates in 4D6 cells expressing (i) GFP-tagged wild-type FOXN1 either alone or (ii) in the presence of untagged Δ550 FOXN1; (iii) quantification of nuclear condensates per cell. (d) Confocal microscopy showing nuclear condensates in 4D6 cells expressing (i) EYFP-tagged Δ550 FOXN1 either alone or (ii) in the presence of untagged wild-type FOXN1; (iii) quantification of nuclear condensates per cell. (e-g) Fluorescence recovery after photobleaching (FRAP) analysis of transfected 4D6 cells. Fluorescence recovery curve of (e) GFP-labelled wild-type FOXN1 and (f) EYFP labelled Δ550 FOXN1 as a function of time. Top: photomicrographs of cell nuclei taken immediately before (0), 6 and 50 seconds (s) after photobleaching of the indicated area (yellow circle). bottom: normalised fluorescence in photobleached area over time; a.u.: arbitrary units. (g) Fluorescence’s recovery rate [arbitrary units (a.u.)/sec] of EYFP-labelled wild-type and Δ550 FOXN1 nuclear condensates following photobleaching. (h). (h) FCS measurement of the diffusion coefficient (µm^2^/sec) of GFP-tagged WT FOXN1 in the nucleus of 4D6 cells within and outside the nuclear condensates and in the presence or absence of untagged Δ550 FOXN1.. Data shown is from 3 (a,c,d,e,f,g) and 2(b) and 1(h) independent experiments. Mean value and SD are shown in panels c,d,g, and h, and the data was analysed using the two-tailed unpaired t-test (c,d,g) and the Kolmogorov-Smirnov test (h). p-values: ≥0.05 (ns),, <0.0001(****). See also Figure S5

We investigated the nuclear behavior of wild-type and Δ550 FOXN1 expressed either alone or in combination. Live imaging of 4D6 cells expressing either wild-type or Δ550 FOXN1 (each labeled separately with a distinct fluorochrome) revealed large nuclear biomolecular condensates resembling aggregates formed by liquid-liquid phase separation and specialized in gene regulation and genome maintenance (Figure 5ci, 5di and Supplementary Figure 5d) [31–36]. The FOXN1 nuclear condensates were also observed in wild-type TEC in thymus tissue sections (Figure 5b) confirming their existence in primary TEC *in situ*. Co-expression of wild-type and Δ550 FOXN1 significantly decreased the number of condensates formed by wild-type FOXN1 (Figure 5cii and ciii) while those containing the Δ550 mutant remained unchanged (Figure 5dii and diii). Thus, Δ550 FOXN1’s ability to interfere with the formation of nuclear condensates is consistent with its disruption of wild-type FOXN1 homo-multimers.

Proteins in membrane-less nuclear organelles characteristically include intrinsically disordered regions (IDRs), whose serine content is required for condensate formation [14–17]. Both wild-type and Δ550 FOXN1 had IDRs with a higher serine content than ordered segments of the proteins (Supplementary Figure 5e and f). The lack of a membrane surrounding nuclear condensates allows rapid exchange of components with the nucleoplasm [12]. We therefore used fluorescence recovery after photobleaching (FRAP) to study movement of fluorescently-labeled FOXN1 in the condensates. Photobleaching of condensates in 4D6 cells expressing either wild-type or Δ550 FOXN1, followed by recovery for 50 seconds (Figure 5e and f), revealed the Δ550 mutant had a reduced recovery rate (Figure 5g), indicating it only partially retains the dynamic properties of wild-type FOXN1. Similarly, FOXN1 variants truncated at either aa 457 or 550 and therefore each with fewer IDRs formed nuclear condensates at a frequency comparable to that of wild-type or Δ550 FOXN1 (Supplementary Figure 5d) but displayed decreased FRAP recovery rates (Supplementary Figure 5g). Hence, different FOXN1 C-terminal mutants could form nuclear condensates but shorter aa sequences significantly impaired exchange of FOXN1 between the condensates and the surrounding nucleoplasm (Figure 5cii).

As transcription factors translocate within the nucleoplasm and bind to their target sequence [37], they can form a range of interactions with other molecules that influence their behavior and function. Within biomolecular condensates, the transcription factor diffusion is slower compared to the rest of the nucleoplasm due to the organelles’ compact nature and the presence of other interacting molecules [31, 33, 38]. Fluorescence correlation spectroscopy (FCS) revealed that diffusion of wild-type FOXN1 was substantially reduced within condensates compared to the adjacent nucleoplasm (11-fold, p<0.0001; ≈0.07 µm^2^/s inside the condensate vs ≈0.8 µm^2^/s outside the condensate; Figure 5h). Diffusion of wild-type FOXN1outside of condensates remained unchanged in the presence of Δ550 FOXN1 (≈0.5 µm^2^/s; Figure 5h). These results are consistent with an exclusion of wild-type FOXN1 from the Δ550 FOXN1 condensates in which the FOXN1 dwell time is typically reduced and correlate with wild-type FOXN1’s reduced transcriptional activity in the presence of Δ550 FOXN1. Thus, wild-type FOXN1 is locally enriched in higher-order liquid-like aggregates, whose diffusion properties change in the presence of Δ550 FOXN1.

### Binding partners modulate FOXN1 transcription

Transcription factors interact with a large number of other molecules that modulate their function. To identify the FOXN1 interactome, we analyzed pull-downs from nuclear lysates of 4D6 cells expressing either wild-type or Δ550 FOXN1. 340 putative binding partners were identified by liquid chromatography with tandem mass spectrometry (LC-MS-MS). To exclude false positives, we required candidate proteins: (i) be detected in all replicates of wild-type or Δ550 FOXN1 precipitates; (ii) not correspond to known contaminants (e.g. albumin, keratins, histones, cytoplasmic proteins and ribosomal proteins) [39]; and (iii) to localize to the nucleus according to the UniProt Knowledgebase [40]. The number of candidates was further reduced to 32 applying two additional suppositions. First, we considered a situation where the quantity of binding partners recovered from precipitates would correlate with the availability of FOXN1 but would be independent of differences in the transcription factors’ C-terminus. Because of the greater expression efficiency of wild-type FOXN1 (Figure 1c and Supplementary Figure 6a), the amount of candidate proteins was expected to be higher in precipitates of wild-type FOXN1 under conditions where the number of captured complexes was not limited (Supplementary Figure 6b). Second, we assumed that a protein partner characteristically associated with wild-type FOXN1 would be sequestered away as a result of a higher affinity for the Δ550 FOXN1 mutant. Here, the amount of the candidate binding partner present in immunoprecipitations would directly correlate with the presence of Δ550 FOXN1 and inversely compare to the concentration associated with wild-type FOXN1 (Supplementary Figure 6b). Of the 32 candidate FOXN1 binding partners, 30 behaved as expected for the first assumption and 2 behaved as expected for the second assumption (Supplementary Table 3). These FOXN1 interactome candidates included (i) mRNA splicing factors, (ii) transcriptional co-activators, (iii) DNA binding and/or (iv) RNA binding proteins (Figure 6a).

**Figure 6:**
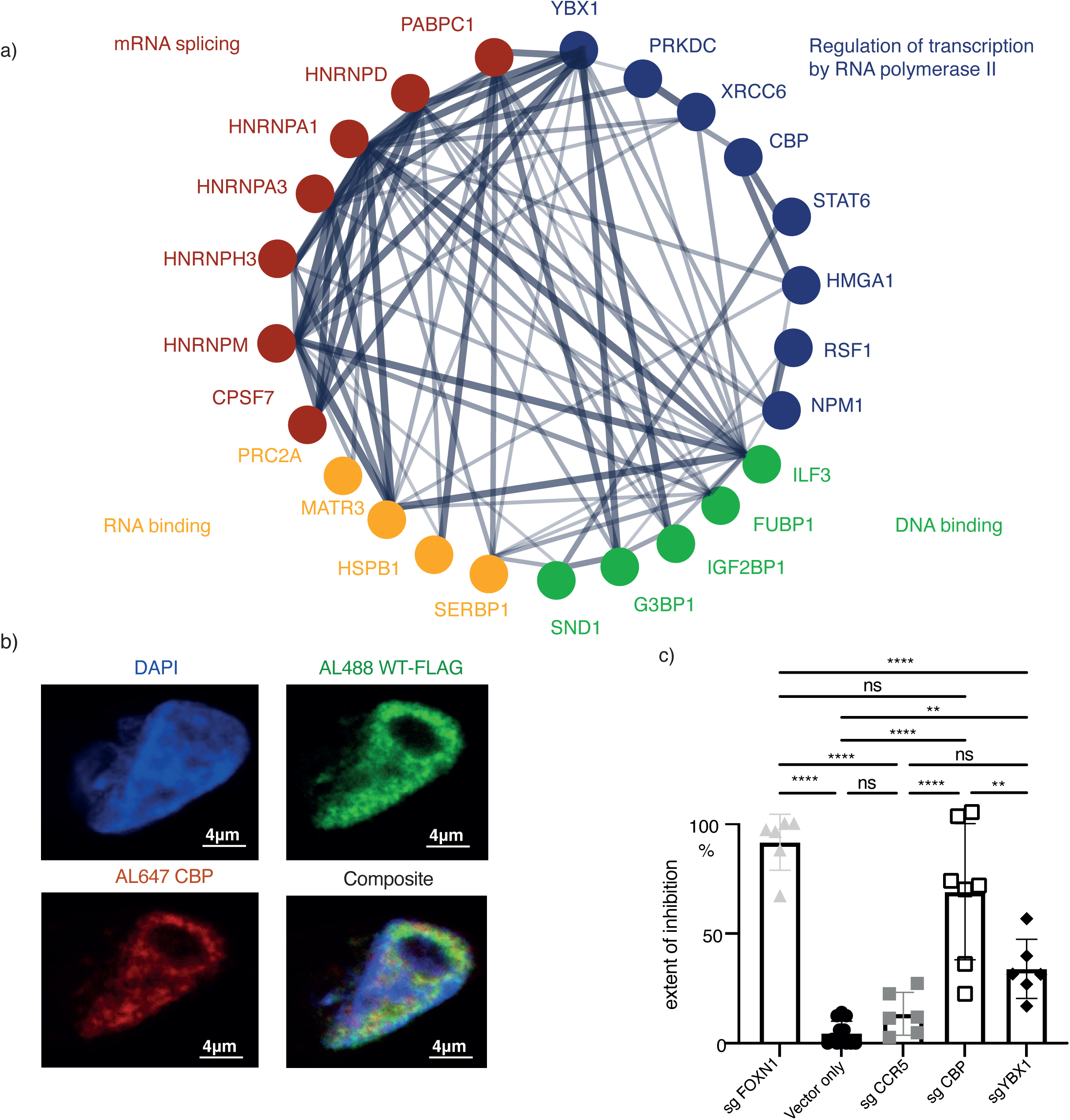
The FOXN1 interactome. (a) STRING protein-protein interaction analysis of FLAG-tagged wild-type and Δ550 FOXN1 pulldowns from transfected 4D6 cells and subsequent analysis by LC-MS-MS. The links show previously verified interactions whereby the thickness of individual lines represents the strength of experimental evidence for such interactions. (b) FOXN1 and CBP co-localize in 4D6 cells. Indirect immunofluorescence microscopy combined with DAPI staining and confocal analysis of FLAG-tagged wild-type (WT) FOXN1 and endogenous CBP in 4D6 cells. The extent of CBP and FOXN1 colocalisation as assessed by Pearson’s correlation equals to 0.54 (where 0 equals to no colocalization and 1 to complete colocalization). (c) CBP and YBX1 are co-factors for FOXN1-mediated transcription. The frequency of cells positive for EYFP was measured using 4D6-EYFP^β5t^ FOXN1^CMV^ cells that had a CrispR-mediated deletion of either CBP, YBX1, FOXN1 or CCR5. 4D6-EYFP^β5t^ FOXN1^CMV^ cells transfected with only the cas9 vector served as an additional control. CrispR-targeted loss of FOXN1 and CCR5 served as a positive and negative control, respectively. The frequency of loss of EYFP expression was calculated relative to cells with loss of FOXN1 expression. The data is representative for at least 2(c) and 1 (b) independent experiments with three technical replicates each. Mean value and SD are shown and statistically compared by using a two-tailed unpaired t-test; p-values: ≥0.05 (ns), 0.0332(*), 0.0002 (***) <0.0001(****). See also Figure S6 and Table S3

Selected interactome candidates were tested for their role in FOXN1-controlled gene transcription using 4D6 cells stably transduced with a lentiviral vector constitutively expressing FOXN1 and containing a minimal *Psmb11* promoter [2] controlling the expression of EYFP (designated 4D6-EYFP^β5t^ FOXN1^CMV^ cells; Supplementary Figure 6c and d). We identified the transcriptional co-activator Creb Binding Protein (CBP) and RNA binding protein Y-box binding protein 1 (YBX1) as potential FOXN1 interaction partners. Both proteins correlated positively with the amount of immunopreciptated FOXN1 protein, contained several IDRs and a higher disordered confidence score (Figure 6a, Supplementary Figure 5f), and could be detected in all TEC subtypes (data not shown [1]. Endogenous CBP also co-localized in transfected 4D6 cells with FOXN1 (Pearson correlation 0.54) (Figure 6b). Deletion of *Cbp* and *Ybx1* resulted in a significant reduction of FOXN1-mediated activation of EYFP in 4D6-EYFP^β5t^FOXN1^CMV^ cells (Figure 6c), demonstrating that both proteins play a role in FOXN1-regulated gene expression.

### FOXN1 expression is developmentally regulated

To understand the mechanism by which heterozygosity for *Δ550 FOXN1* and *Δ505 FOXN1* could initially support thymus organogenesis but over time give way to thymic aplasia or hypoplasia, we reasoned that given the dominant negative effect of these mutants was incomplete (Figure 4b and Supplementary Figure 7a), sufficient wild-type FOXN1 activity could be generated from a highly active *FOXN1* promoter during early phases of thymus organogenesis. To test this idea, we engineered mice that expressed a fluorescent timer protein (FTP) under the control of the *FOXN1* promoter. The FTP initially emits a blue fluorescence (peak at 465 nm) but progressively matures within 24 hr to emit a red fluorescence (604 nm; Supplementary Figure 7b) [41]. The FTP’s geometric mean fluorescent intensity (gMFI) at 465 nm correlated with *FOXN1* promoter strength. gFMI gradually increased from E12.5, reaching a plateau 4 days later until birth, when it rapidly diminished to values below those observed at early embryonic time points (Figure 7a and Supplementary Figure 7c). These dynamic shifts were paralleled by changes in FOXN1 protein levels (Figure 7b). Thus, the activity of the FOXN1 promoter varies significantly over time. This, together with the observation that heterozygosity for the Δ505 allele diminished FOXN1 protein levels in TEC supports our hypothesis that residual mouse FOXN1 activity is insufficient for normal thymus function in FOXN1^WT/Δ505^ mice after birth.

**Figure 7:**
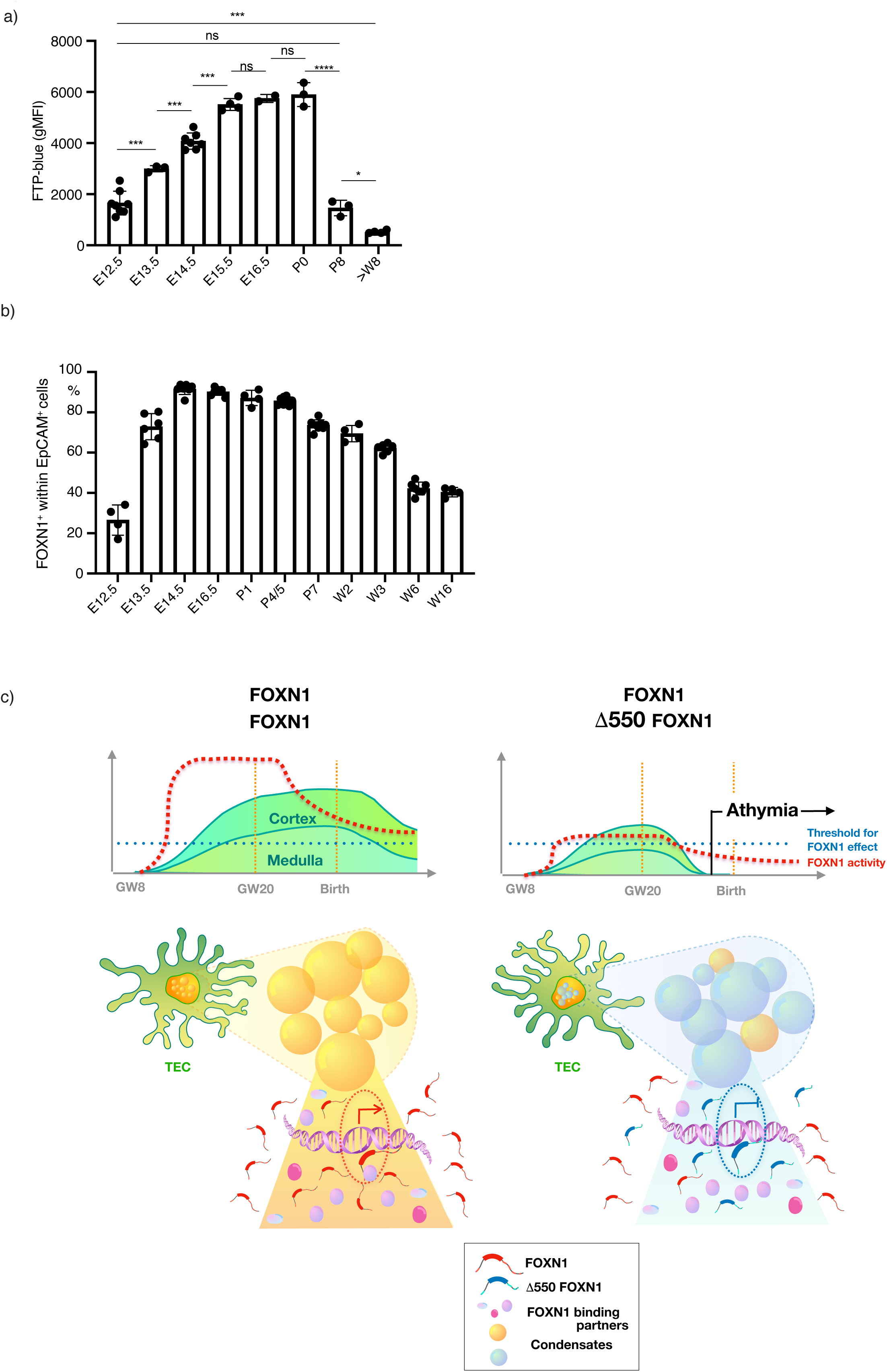
*FOXN1* promoter activity is developmentally regulated. (a) Geometric mean fluorescent intensity (gMFI) of fluorescent timer protein at 465 nm (FTP-blue) in total TEC from FTP^FOXN1^mice at indicated ages. Each dot represents an individual mouse (b) frequency of FOXN1^+^ cells in total TEC from FOXN1^WT/WT^ mice at indicated ages. The data is from 2 independent experiments at each time point with at least three mice each. Mean value and SD are shown in bar graphs and were calculated with two-tailed unpaired t-test; 0.05 (ns), 0.0332(*), 0.0021(**), 0.0002 (***) See also Figure S7 (c) Graphical abstract of the model explaining how Δ550 FOXN1 compromises thymus tissue maintenance. Nuclear FOXN1 (red symbols) concentration is required to be above a critical limit (blue dotted line) to initiate and maintain TEC differentiation. Owing to its dynamic promoter activity, the level of FOXN1 (red broken line) greatly exceeds this threshold during the second trimester and remains sufficiently elevated for an extended period of time thereafter. This allows FOXN1 to accumulate in sufficient quantities in nuclear condensates (yellow spheres), interact there with essential binding partners and drive the transcription of target genes (left). In individuals heterozygous for the Δ550 FOXN1 mutant (blue symbols), wild-type FOXN1 is however expressed from a single allele and its concentration is thus reduced (right). Furthermore, wild-type FOXN1 molecules are competitively displaced by Δ550 FOXN1 from condensates so that the majority of these structures contain mostly the transcriptionally inactive variant (blue spheres). After the second trimester transcription of the *FOXN1* locus is reduced and consequently only few condensates retain wild-type FOXN1-controlled transcriptional activity, which collectively are insufficient to secure FOXN1 activity above its critical threshold. As a consequence, TEC are not maintained and functional thymus tissue is lost causing athymia with peripheral T lymphopenia.

## Discussion

FOXN1 is a gene dosage sensitive transcription factor required for the differentiation and function of thymic epithelia. Mutations in FOXN1 have so far been linked to a recessive loss-of-function and more recently to hypomorphic variants believed to cause pathology through reduced gene dosage [4, 8, 10, 21, 42]. Individuals heterozygous for *FOXN1* mutations that alter the protein’s C-terminus are characteristically T-cell lymphopenic (primarily affecting CD8 T cells) and, when symptomatic, display various pathologies including severe recurrent infections [21, 42]. Here we describe a spontaneous *FOXN1* variant and elucidate the mechanism by which this and other C-terminal mutants exert a dominant negative effect on wild-type FOXN1.

Our data reveal that wild-type FOXN1 forms multimers that interact with other nuclear factors to form condensates poised to initiate transcription. FOXN1 condensate formation and DNA binding via its forkhead (FKH) domain require sequences in the C-terminal domain, whereas the N-terminal domain is dispensable for DNA binding [10]. Nuclear condensates mediate transcription by concentrating and promoting efficient molecular interactions within a discrete biochemical milieu [31–34]. Immunoprecipitations with wild-type FOXN1 identified several putative interacting partners commonly detected in these nuclear structures and collectively involved in mRNA splicing, transcriptional regulation, and DNA/RNA binding [31, 33, 43, 44]. One such partner is the transcriptional co-activator CBP, which serves as a central node in eukaryotic transcriptional networks by interacting via its IDRs with other proteins [17, 45]. IDRs, which are also present in FOXN1 and YBX1, use a small number of residues [16] or their post-translational modifications [17] to promote high-specificity, modest-affinity interactions [46, 47]. The role of FOXN1’s interaction with CBP and, to a lesser extent, YBXI, is demonstrated by the requirement of both co-factors to initiate *Psmb11* promoter-controlled gene transcription. FOXN1’s binding to CBP is unlikely to be critical for inclusion in condensates as the Δ550 mutant fails to bind the co-factor but is still contained within these structures.

The shortened Δ550 FOXN1 with its partially altered C-terminal sequence preserves 3 of the 6 IDRs present in wild-type FOXN1 and retains the capacity to form condensates. Although IDRs have been suggested to enable phase separation, this capacity may, alternatively, be controlled by sequences with negative charges, high number of aromatic/hydrophobic residues and/or a significant serine bias [16] [14]. Independent of the precise mechanism in play, the difference in sequence between wild-type and mutant FOXN1 demonstrate that the three terminal IDRs are not required for condensate formation. Instead, changes to the C-terminal sequence correlate with increased residence, separating condensate formation from dwell time.

Our *in vitro* experiments indicate that the DNA sequences bound by Δ550 FOXN1 are in part different compared to those bound by wild-type FOXN1, suggesting that the C-terminal sequences influence FOXN1 interactions with proximal gene regulatory regions. Nuclear condensate formation by the mutant FOXN1 is, however, unimpaired, indicating that condensate formation does not require CBP binding, although CBP is required for wild-type FOXN1 transcriptional activity. The changes in the C-terminal sequence increase Δ550 FOXN1’s dwell time within nuclear condensates. Thus, in cells expressing both wild-type and Δ550 FOXN1, the latter preferentially accumulates in the condensates due to its longer residence time, displacing wild-type FOXN1 and precluding it from activating transcription. It is also possible that the lack of CBP binding to Δ550 FOXN1 impairs normal displacement of the transcriptionally inactive interactome from DNA binding, further blocking access of wild-type FOXN1 to its target sequences. Functional FOXN1 is not completely excluded from nuclear condensates in the presence of Δ550 FOXN1, as limited gene transcription by wild-type FOXN1 is seen under co-expression conditions.

The heterozygous expression of Δ505 *Foxn1* in mice, the orthologue of the human mutation, results in thymic hypoplasia and changes in the composition of the TEC scaffold. The frequency and absolute cellularity of cortical epithelia is increased in FOXN1^WT/Δ505^ mice, whereas in wild-type mice perinatal cTEC represent approximately one-third of all thymic epithelia at week 1 of age but three weeks later only contribute less than 1% of the epithelial scaffold[1]. The differentiation of perinatal cTEC to mature epithelia is, however, unaffected in the presence of Δ505 FOXN1 consistent with the idea that a decrease in the normal FOXN1 gene dosage still allows for the maintenance of perinatal cTEC and is sufficient for their progression to mature cortical TEC.

TEC cellularity is unchanged in FOXN1^WT/Δ505^ animals despite thymic hypoplasia, underscoring that the functional competence of the TEC scaffold is compromised by the expression of the mutant FOXN1 and the consequent changes in the frequency of individual TEC subtypes. The frequency of thymocytes undergoing programmed cell death is reduced among CD4+ CD8+ cells indicating a qualitative deficiency of mutant cTEC when compared to wild-type epithelia. In contrast to cTEC, almost all other TEC subtypes in FOXN1^WT/Δ505^ animals are reduced in their frequency. Moreover, mTEC display a partial block in lineage maturation, reduced MHC class II expression, and impaired expression of peripheral tissue specific antigens, a phenomenon known as promiscuous gene expression and important to establish central T cell tolerance [25]. These changes parallel a reduction in late maturational stages of post selection thymocytes. Moreover, the frequency of NKT cells, but not that of Treg, is reduced in FOXN1^WT/Δ505^ mice. As both lymphoid lineages depend on specific TEC niches, differential changes in the epithelial scaffold of FOXN1^WT/Δ505^ mice must account for this variance. This conclusion is consistent with expression of a dominant negative form of FOXN1 affecting TEC maturation differently depending on the cells’ lineage and developmental stage.

Finally, our study indicates why individuals heterozygous for the Δ550 FOXN1 mutation are athymic at birth, and yet were able to produce a limited population of peripheral T cells. Our data suggests that physiological changes in *FOXN1* promoter activity account for the observed loss of thymic tissue during the later stages of gestation (Figure 7c). Early in development high levels of wild-type FOXN1 are generated due to the high activity of the *FOXN1*promoter and this provides sufficient wild-type protein to mediate transcription despite the presence of the dominant negative mutant variant. This balance in favor of wild-type FOXN1 is further helped by the mRNA encoding Δ550 FOXN1 being less efficiently translated when compared to wild-type transcripts. With increasing age, *FOXN1* promoter activity decreases and the availability of wild-type FOXN1 protein is reduced to a nuclear concentration below the critical threshold required to maintain TEC function [2, 8]. The exact time when thymopoiesis stops in *Δ550 FOXN1* heterozygous patients can be estimated to have happened after week 20 of gestation because N-nucleotide additions to the TCR beta chain’s CDR3 are similar for the index patient and healthy individuals, arguing for a time point when the completion of thymus organogenesis and the expression of terminal deoxynucleotidyl transferase (TdT) has already occurred [48] [49]. In the Δ505 FOXN1 mouse model, this time point corresponds to the first days after birth[50] and is consistent with thymus hypoplasia being observed in these animals only after birth.

Taken together we identify a critical mechanistic determinant of thymic development by demonstrating that FOXN1 participates in the formation of nuclear condensates essential for transcription. Moreover, we provide a molecular explanation how a FOXN1 mutant dominantly interferes with the dynamic formation of transcriptional hubs ultimately causing a novel form of a primary immunodeficiency. The concept that liquid-liquid phase separation in biomolecular condensates helps compartmentalize distinct cellular processes has recently been discerned. Here, we unequivocally demonstrate a biological role for nuclear condensates in the function of a key developmental transcription factor as well as showing that spontaneously occurring variants of this factor act to cause disease by interfering with condensate formation.

## Acknowledgements

We thank Raif Geha, and Thomas Barthlott for helpful discussions and Guy Riddihough (LSE) for editorial assistance. We thank Sam Palmer for generating the graph in Supplementary Figure 5g. This study was supported by grants from the Swiss National Science Foundation (IZLJZ3_171050; 310030_184672) and the Wellcome Trust (105045/Z/14/Z)) to G.A.H. G.A.H. was also supported by the National Institute for Health Research (NIHR) Oxford Biomedical Research Centre (BRC). I.A.R was a recipient of a Departmental stipend (Department of Paediatrics, University of Oxford). A.H was supported by an NIHR Clinical Lectureship. FD was supported by the Wellcome Trust [109032/Z/15/Z] and an NIHR Clinical Lectureship. S.C. was a recipient of the international postdoc fellowship from the Swedish research council (Vetenskapsrådet). FK is a recipient of the Swiss National Science Foundation - Early Postdoc.Mobility Fellowship - P2BSP3_188183. ES is supported by SciLifeLab Fellowship and Swedish Research Council Starting Grant (2020-02682). The OxClinWGS study was supported in part by the Wellcome Trust and Department of Health as part of a Health Innovation Challenge Fund scheme grant (Wellcome Core Award Grant Number 203141/Z/16/Z) to J.C.T. J.C.T, A.T.P and C.C. were also supported by the National Institute for Health Research (NIHR) Oxford Biomedical Research Centre (BRC). The views expressed are those of the authors and not necessarily those of the NHS, the NIHR or the Department of Health or Wellcome Trust. GA is support by an MRC Programme Grant (MR/T029765/1). WQ, EGD and ASG were supported by the NIHR (RP-2014-05-007) and the Great Ormond Street Hospital Biomedical Research Centre.

## Author contributions

G.A.H supervised and conceptualized the project; I.A.R, A.H, G.A.H designed the experiments; I.A.R, S.M.,F.K, S.C, F.D, NP and M.E.D performed the mouse experiments and/or analysed and interpreted the results; A.H performed all the ChIP-Sequencing, single cell and bulk RNA sequencing analysis and interpreted the results; I.A.R performed the in-vitro experiments and analysed the results; I.A.R, M.E.D, E.A designed and generated the tagged FOXN1 constructs; I.A.R, E.S, F.K, C.L, S.Z, performed and/or analysed and interpreted the confocal microscopy experiments; P.D.C analysed the proteomics data; P.H designed and generated the FOXN1^WT/Δ505^ mice; A.W, K.J, provided the thymus from FOXN1^WT/nude^ mice and antibody reagents; FD, WQ, GD, CH identified, recruited and phenotyped the patients, A.P, C.C, F.D, H.D., EA and J.T performed the Whole Genome Sequencing analysis and/or interpreted the results; J,A, N and O.G, designed and performed the EMSA assay and interpreted the results; Y.S.M and T.F. assisted in the design and execution of the CrispR KO experiments, A.S.G performed the TCR MiSEQ analysis; G.A provided the FOXN1^WT/nude^ mice, CD1d tetramer and contributed to data interpretation. G.A.H, I.A.R and A.H wrote the manuscript and prepared the figures.

## MATERIALS AND METHODS

### Patient Recruitment

The patient was recruited as part of the OxClinWGS programme to apply clinical-grade whole genome sequencing for patients with a broad range of rare diseases to identify the pathogenic mutations. Written informed consent was obtained from the patient as part of a study approved by the South Central Research Ethics Committee reference REC 12/SC/0044.

### Whole genome sequencing analysis pipeline

DNA was extracted from peripheral blood mononuclear cells (PBMCs) of the father, mother sibling and the index patient using a magnetic bead method and the QIAsymphony automated extraction system (Qiagen). TruSeq PCR Free libraries were prepared for all samples. Genome Sequencing was performed with the HiSeq2500 (Illumina) instrument at the clinically accredited Molecular Diagnostics Laboratory at the John Radcliffe Hospital. Paired 100bp reads were mapped to hs37d5 using BWA v.0.7.10 and Stampy v1.0.23 (www.well.ox.ac.uk/stampy). Samtools v1.3 was used to sort and merge the BAM files and Picard markduplicates v1.111 was used to remove duplicates from the BAM files. The mean read depth was 30X. Variant calling was performed with Platypus (www.well.ox.ac.uk/platypus) using default settings except for minFlank=0.

Variants were filtered for those with a call quality of at least 20 that passed upstream pipeline filtering. Variants also were required to have an allele fraction at least 5% and lie outside the top 5% most exonically variable 100bp windows in healthy public genomes (1000 genomes). Variants also had to lie outside the top 1% most exonically variable genes in healthy public genomes (1000 genomes). We then excluded variants that are observed with a population allele frequency of ≥ 0.1% in the 1000 genomes project or ≥ 0.1% in the NHLBI ESP exomes (all) or ≥ 0.1% in the AFC or ≥ 0.1% in ExAC (maximum) or ≥ 0.1% in gnomAD (maximum). We then kept exonic variants (up to 20bp into the intron) that are frameshifts, in-frame indels, stop codon changes, missense, or that splice site-altering variants (within 8bp into intron or predicted to disrupt splicing by MaxEntScan). Lastly, we filtered for variants which were heterozygous in all 3 affected family members and which were not seen in the unaffected mother.

Variant filtering was performed using QIAGEN Clinical Insight Interpret version 7.1.20210316 (https://variants.ingenuity.com/qci) and data was exported on 17^th^ March 2021. The content versions of databases used in this commercial software are as follows: CADD (v1.6), Allele Frequency Community (2019-09-25), EVS (ESP6500SI-V2), Refseq Gene Model (2020-04-06), JASPAR (2013-11), Ingenuity Knowledge Base Snapshot Timestamp (2021-03-02 01:49:28.442), Vista Enhancer (2012-07), Clinical Trials (B-release), MITOMAP: A Human Mitochondrial Genome Database. www.mitomap.org, 2019 (2020-06-19), PolyPhen-2 (v2.2.2), 1000 Genome Frequency (phase3v5b), ExAC (0.3.1), iva (Nov 20), TargetScan (7.2), phyloP (GRCh37/hg19) 2014-02), GENCODE (Release 33), CentoMD (5.3), Ingenuity Knowledge Base (B-release), OMIM (July 06, 2020), gnomAD (2.1.1), BSIFT (2016-02-23), TCGA (2013-09-05), ClinVar (2020-09-15), DGV (2016-05-15), HGMD (2020.4), dbSNP153 (GRCh37/hg19), SIFT4G (2016-02-23).

### Cell culture and transfection

4D6 cells [53] were grown in RPMI 1640 (Sigma-Aldrich) supplemented with 10% (v/v) heat inactivated fetal bovine serum (FBS) (SLS), 10 mM HEPES buffer, 1% (v/v) non-essential amino acids (NEAA) (Lonza), 2mM L-glutamine (Lonza), 100U/ml penicillin, and 100µg/ml streptomycin (Lonza) at 37°C with 5% CO_2_ in a humidified incubator. Murine TEC1.2 cells were grown in IMDM (Life Technologies) supplemented with 2% (v/v) heat inactivated FBS (SLS), 38.6mM NaHCO_3_ (Sigma), 50mM 2-Mercaptoethanol (Sigma) and 0.3g/L Primatone RL/UF (Fluka), at 37°C with 10% CO_2_ in a humidified incubator. Cells were seeded in cell culture plates of various sizes (Corning) and allowed to grow to reach 70% confluency by the time of transfection. Within 18-20 hours after seeding, cells were transfected with plasmids of interest using Fugene (Promega) as per the manufacturer’s recommendations. Four hours post transfection, the cell culture media was replaced by fresh media. Cells were harvested 24- and 48-post transfection for further analysis.

### Plasmid construction

Human FOXN1 cDNA was purchased from Genecopoeia (sequence accession number BC140423). Full length FOXN1 cDNA without a stop codon was cloned into vector backbones positioning at its C-terminus either a flag (pCSF107mT-GATEWAY-3’-FLAG, Addgene), myc (pCSF107mT-GATEWAY-3’-Myc tag, Addgene), green fluorescence protein (GFP) or an enhanced yellow fluorescent protein (EYFP) sequence (pcDNA1.3-, Addgene) using standard cloning methods. Mouse wild-type FOXN1 was cloned in a pLKP1 vector and Δ550 FOXN1 flag in a MR207953 vector using standard cloning methods. Insertion of single base pair changes for the generation of human and mouse FOXN1 variants was achieved by site-directed mutagenesis using the Phusion site-directed mutagenesis kit (Thermo Fisher Scientific).Specific primers for the introduction of single base pair deletions or single base pair changes were designed using the following software www.agilent.com/store/primerDesignProgram.jsp. All plasmids were verified by Sanger sequencing.

### RNA, cDNA synthesis and qRT-PCR

RNA isolation was performed using the Rneasy Mini (Qiagen) or Rneasy Plus Micro kit (Qiagen). cDNA was synthesized using SensiFAST cDNA kit (Bioline) and assessed by qPCR (SensiMix SYBR Hi-Rox, Bioline). Expression levels of each gene were normalized against *GAPDH* expression and fold change was calculated using the 2^-ΔCT^ or 2^-ΔΔCT^ equations [54]. The following primer sequences were used: human *GAPDH*: F: 5′-AACAGCCTCAAGATCATCAGC-3, R: 5′-CTGTTGCTGTAGCCAAATTCG-3′; human *FOXN1*: F: 5′-TTCCTTACTTCAAGACAGCAC-3′ R: 5′-GGTTCTTGCCAGGAATGG-3′ mouse *Gapdh*: F: 5’GGTGAAGGTCGGTGTGAACG3’, R: 5’ACCATGTAGTTGAGGTCAATGAAGG3’, mouse *Foxn1*: F: 5’TCTACAATTTCATGACGGAGC3’ R: 5’TCCACCTTCTCAAAGCACTTGTT3’

### Luciferase reporter assay

The luciferase reporter assay was performed as described previously [2]. In short, 4D6 cells were co-transfected with a renilla control plasmid (pRL Promega), and a luciferase reporter plasmid (pGL4.10[Luc2], Promega) under the control of a minimal wild-type *Psmb11* promoter (designated β5t-luc) or a β5t promoter with a mutated FOXN1 binding sites (β5t-mut-luc) plus a FOXN1 construct of interest in a ratio of 1:10:10. Luciferease activity was measured 24 h later using the Dual-Lucifearase Assay kit (Promega).

### Protein preparations and Western blot

For Western blot analysis, 4D6 were treated with lysis buffer [25mM Tris-HCL pH7.5] 50mM NaCl (Sigma), 0.1%NP-40 (Sigma), 0.1% SDS (Sigma), 0.5% Sodium deoxycholate (Sigma), 10%Glycerol (Sigma), and protease inhibitors (1 tablet/10ml, Roche)] followed by SDS-PAGE and immunoblotting, using mouse anti-FLAG (monoclonal M2, Merck), rabbit anti-myc (Cell Signaling) and rabbit anti-GAPDH (Cell signaling) antibodies. For the relative quantification of FOXN1 in figure 1c, the loading control was run through the gel from the same loading well as the experimental sample. After the protein transfer, the membrane was cut in two parts at about 50KD and each part was developed separately with anti-FLAG (top part) and anti-GAPDH (bottom part). For the relative quantification of FOXN1 protein levels, the fluorescence intensity of WT or Δ550 mutant FOXN1 protein bands were calculated relative to the fluorescence intensity of the GAPDH loading control after subtracting background fluorescence.

### Immunofluorescence

4D6 cells were plated into 12 well cell culture plates containing sterile coverslips coated with lysine (1:10 dilution in PBS, Sigma) and transfected as described earlier. For the immunofluorescent staining cells were fixed with 4% paraformaldehyde solution and permeabilized with 0.2% Triton X-100, then blocked in blocking buffer [1x PBS, 1% BSA, 5% goat serum, (all Sigma reagents) and subsequently stained with primary antibodies: Mouse anti-FLAG (1:500) (monoclonal M2, Merck) or rabbit anti-myc (1:200) (monoclonal, Cell Signaling) and secondary antibodies [goat anti rabbit Alexa Fluor 488, goat anti-mouse Alexa Fluor 647,(Sigma)]. Where indicated, 4D6 cells were counter-stained with DAPI (10pg/mL in methanol) and all slides were mounted with ProLong Gold Antifade Mountant (ThermoFisher Scientific). A ZeissLSM 880 inverted confocal laser scanning microscope was used to visualize the stained cells.

Frozen tissue sections were cut at 7µm thickness, fixed with 3.6% formalin, permeabilized with 0.2% Triton X-100 and stained with R-Phycoerythrin-labeled (Abcam) anti-FOXN1 (a generous gift of HR Rodewald)[24], anti-deltaNp63 (Biolegend) and anti-AIRE (Invitrogen) antibodies. Unlabeled primary antibodies were detected using secondary antibodies (goat anti-rat Alexa Fluor 488 or goat anti-rabbit Alexa Fluor 647; both from Thermo Fisher Scientific). Tissue was counterstained with DAPI (10pg/ml final dilution in PBS) and mounted with Hydromount Histology mounting media (National Diagnostics). Images were acquired using a Leica DMi8 microscope and analysed by Fiji (ImageJ software).

### RNA-sequencing

4D6 cells were transfected with plasmids encoding 1) wild-type-FOXN1 alone 2) *Δ550*-FOXN1 mutant alone, 3) wild-type and Δ550 οr 4) empty vector only, along with a GFP-expressing vector in a 4:1 ratio to facilitate sorting of positively transfected cells. Three biological replicates were prepared for each sample. A total of 75,000 GFP-positive cells were FACS sorted and RNA was extracted using the RNeasy Plus Micro kit (Qiagen). RNA was processed using the RNA-Seq Poly A method and 100 bp paired-end RNA-Seq was performed on the Illumina HiSeq4000 platform (Wellcome Centre for Human Genetics, University of Oxford).

For data analysis, reads were aligned against the Ensembl transcriptome and genome (GRCh38.EBVB95-8wt.ERCC) using STAR (version 2.5.3a). The allocation of reads to protein coding gene meta-features was done using HTSeq. Differential expression analysis on genes, with at least one aligned fragment, was performed using general linear modelling in edgeR with trimmed-M-of-means library size correction and tagwise dispersion estimates. Differentially expressed genes were identified using the default edgeR threshold (FDR<0.05). Gene ontology analysis was performed using ClusterProfiler. The third biological replicate of the *Δ550* sample was removed as an outlier due to a low number of detected genes and initial exploratory analysis using multi-dimensional scaling of inter-sample differences.

### Single-cell RNA-sequencing of FACs sorted TEC

Single cell libraries were processed using Cellranger (version 3.1.0). Cells were excluded from analysis if the total UMI count was <1,000 or >32,000, the number of detectable features was <562 or >7,943 or the proportion of UMIs mapping to mitochondrial genes was >10%. Cells from individual 10X Chromium lanes were combined using canonical correlation analysis-based integration in Seurat (version 3.1.1)[55]. Cell clusters were identified using the default resolution of 0.8 and visualized using UMAP Cell type annotation was based on projection from our previously published reference data [1]along with manual annotation of non-TEC clusters. Differential analysis was conducted in Seurat using FindMarkers with default settings. ClusterProfiler was used for gene ontology analysis[56].

### Electrophoretic Mobility Shift Assay (EMSA)

EMSA was performed as recently described [9], using purified wild-type FOXN1 and *Δ550* FOXN1 constructs. The sequence of the DNA probe used was part of the minimal promoter region of the proteasome subunit alpha 7 (PSMA7), a high-confidence FOXN1 target [2] with the following sequence 5’GCAGCA**GACGC**AACAGAGCGA**GACGC**CAGGG3’ (with FOXN1 consensus sites in bold).

### Chromatin Immunoprecipitation (ChIP) followed by DNA-Sequencing

4D6 cells were transfected with a construct encoding either wild-type or the Δ550 FOXN1 mutant and tagged with a FLAG sequence. Three biological replicates of each of these two conditions were collected for ChIP-Seq analyses. Chromatin immunoprecipitation was performed using the iDeal ChIP-seq kit for Transcription Factors (Diagenode) as per the manufacturer’s recommendations. Briefly, 2.5 x10^6^ transfected cells were collected and subjected to protein-DNA crosslinking for 20 min. Cells were then lysed and chromatin was sheared for 8 sonication cycles (30’’ ON/30’’ OFF). Sheared chromatin was immunoprecipitated using M2 anti-FLAG antibody (Sigma) while parallel input samples from non-immunoprecipitated chromatin were prepared for each biological replicate of each condition. DNA libraries for all 12 samples were prepared using the MicroPlex Library Preparation Kit (12 indexes; Diagenode) following the manufacturer’s recommendations. Libraries were pooled and sequenced at the Wellcome Centre for Human Genetics, University of Oxford, using 75 bp paired-end sequencing emplying an Illumina HiSeq4000 platform. For data analysis, contaminating adaptor sequences were removed from fastq sequences using Trimmomatic (version 0.32). Reads were aligned against the human genome (GRCh38.EBVB95-8wt.ERCC) using STAR (version 2.5.3a). Peaks were called on deduplicated aligned sequences (paired ChIP and input samples) using MACS2 (version 2.0.10) with a relaxed p value setting of 0.1. Peaks from replicates were then pooled and analyzed using irreproducible discovery rate analysis (IDR < 0.05) (https://sites.google.com/site/anshulkundaje/projects/idr). Peaks were filtered against the ENCODE blacklist regions (https://sites.google.com/site/anshulkundaje/projects/blacklists). *De novo* motif identification was performed using MEME-ChIP [29]

### Immunoprecipitation of tagged FOXN1 variants

4D6 cells were transfected and co-transfected to express either FLAG labeled wild-type, FLAG labeled Δ550-FLAG, or FLAG and myc labelled wild-type FOXN1 in the presence or absence of unlabelled Δ550 FOXN1. Cell lysates were incubated with anti-FLAG or anti-Myc antibodies in the presence of protein G or A magnetic beads (Thermofisher Scientific) for 2 hours at 4°C while slowly rotating. For pull downs to be analyzed by Western blotting, beads were boiled at 95°C with denaturating sample buffer [250 mM Tris-HCl [pH 6.8], 10% [w/v] SDS, 35% [v/v] glycerol, 0.05% BPB, and 0.7 M 2-mercaptoethanol]. For pull downs to be analysed by mass spectrometry, beads were incubated with DSSO (disuccinimidyl sulfoxide; Sigma) at a final concentration of 1mM at room temperature for 45min with shaking according to the manufacturer’s recommendations. The reaction was paused by adding 10ul of Tris-HCL ph7.5 1M for 5min and beads were washed 3x with PBS. Proteins were eluted from beads using 40 µg of 3x FLAG peptide (Merck) for 200µL of IP sample incubated on a rotator for 45 min at 4°C.

### Live cell imaging

4D6 cells were plated in 35 mm glass bottom dishes (Ibidi) in the 4D6 cell growth medium described earlier and were then transfected with various combinations of the wild-type and mutant FOXN1 constructs. Twenty-four hours after transfection the culture medium was exchanged with Leibovitz 15 medium lacking phenol red (Thermo Fisher Scientific) and cells were imaged with a ZeissLSM 880 inverted confocal laser scanning microscope. A laser wavelength of 488 nm wave length was used to excite GFP or EYFP. Laser power was kept 50 µW or below to avoid phototoxicity.

### Fluorescence recovery after photobleaching (FRAP)

FRAP was performed in defined areas of the nucleus of 4D6 cells transfected with wild-type, Δ550, Δ457 or Δ550b FOXN1 variants tagged with EYFP using a LSM 880 inverted confocal laser scanning microscope with 40X W (1.2) objective. Fluorescence recovery within the bleached area was normalized to the fluorescence within a reference area outside the bleached area. To calculate the recovery rate, normalized fluorescence recovery was further normalized to the mean pre-bleached intensity. Normalized data was fitted using the curve fitting function in Fiji. The following fitting formula was used: F(t)=a•(1-exp(-b•t)) + F(0), where “a” is the fractional amplitude of a slowly recovering fraction, “b” is the recovery rate, and “F(0)” is the fractional fluorescence intensity at time zero. [57].

### Fluorescence correlation spectroscopy of wild-type and *Δ550* FOXN1

A LSM 780 inverted confocal laser scanning microscope with 40X W (1.2) objective was used. The microscope was calibrated prior to acquiring the data and the point spread function was determined using 10 nM Alexa488 solution. Fluorescence correlation spectroscopy on cells was performed in defined areas of the nucleus of 4D6 cells transfected with wild-type-FOXN1-GFP.. A fluorescence image was first taken of the cell’s nuclei containing fluorescently labelled FOXN1, subsequently, the nuclear location for the FCS analysis was selected and changes in fluorescence were measured at this position as a function of time. For each selected position five FCS measurements of 5 sec each were taken. FCS curves from single runs were individually sorted to determine whether they met the designated criteria which stipulated photobleaching during acquisition time is less than 10% [58]. Curves that did not meet these criteria were excluded from further analysis. Selected FCS curves from each run were combined and fitted using the FoCuS-point software with a 3D and triplet model[59].

### Mass spectrometry proteomics analysis

4D6 cells were transfected with constructs encoding either wild-type-FLAG or Δ550-FLAG sequences. Three experimental replicates were prepared for each experimental condition. Cells were lysed and anti-FLAG immunoprecipitation was performed as described above. To confirm the presence of proteins other than FOXN1, a fraction of the sample was analysed by silver staining (Pierce^TM^ silver staining kit, Thermo Fisher Scientific) according to the manufacturer’s recommendations. For the mass spectrometric analysis, frozen samples were thawed and subjected to two rounds of chloroform/methanol precipitation. An in-solution trypsin digestion was performed as previously described [60]. After digestion and desalting, samples were resuspended in 20µL 2% acetonitrile, 0.1% formic acid and subjected to nano-flow liquid chromatography tandem mass spectrometry (LC-MS/MS, here referred to as MS) [60]. LC MS/MS analysis was performed on an Orbitrap Fusion Lumos instrument (Thermo Fisher Scientific) coupled to an Ultimate 3000 nUPLC with an EASY-Spray column (50 cm). Peptides were separated using a solvent gradient of 2-35% Acetonitrile in 0.1% formic acid/5% DMSO over 60 minutes. MS data was acquired using a method to also cater for DSSO crosslinked peptides [61]. Briefly, precursor masses were acquired with a resolution of 60000 for up to 50 ms. MS2 were acquired in the Orbitrap after CID fragmentation. A mass difference of 31.9721 triggered subsequent MS3 scans with increased collision energy (25% à 35% and detection in the linear ion trap in rapid mode followed by an MS2 scan after ETD fragmentation and detection in the Orbitrap with a resolution of 15000.. The raw MS data were processed with Proteome Discoverer 3.5 including the XLinkX node ([62]) and Progenesis QI for Proteomics v. 4.1.6675.48614 to generate label-free relative quantitation after protein identification in Mascot 2.5 Matrix Sciences. For this, MS/MS data was searched against a swissprot database (retrieved 05/2018) with precursor mass tolerance of 10 ppm and fragment mass tolerance of 0.5Da. Peptide false discovery rate was adjusted to 1% and peptides with Mascot scores <20 removed, before importing the data into Progenesis QI. Detected crosslinks were not followed up.The raw quantitation values were further analysed in R version 3.5.5. For each protein, correlation with FOXN1 was assessed by Spearman’s ⍴, and associated p-values were calculated by the algorithm AS89 [63]. False discovery rate (FDR) estimates at each p-value cut-off (q-values) were calculated from the list of p-values using the fdrtool package (R package version 1.2.15. (https://CRAN.Rproject.org/package=fdrtool) [64]. An FDR cut-off of 5% was applied to data analysis. The STRING protein-protein interaction database, i.e. a database of known and predicted protein–protein interactions containing information from experimental data, computational predictions and public text collections https://string-db.org, was queried to construct a potential interaction network. The accession numbers of all proteins that were detected in all replicates of wild-type or Δ550 FOXN1 precipitates, that did not correspond to known contaminants (e.g albumins, keratins, histones, cytoplasmic proteins and ribosomal proteins) [39] and that localized to the nucleus according to the UniProt Knowledgebase [40] were submitted to the STRING database.

### Generation of the EYFP^β5t^FOXN1^CMV^ 4D6 reporter cell line

4D6 cells were transduced with the in-house designed lentivector EYFP^β5t^FOXN1^CMV^ that was commercially produced (Vector Builder). Stable integration of this lentivector into the genome of 4D6 cells allowed the constitutive expression of FOXN1 under the control of the CMV promoter. Functional FOXN1 bound to the β5t promoter (the same promoter sequence used for the luciferase assays described above) controlled the expression of EYFP.

For the production of infectious lentivirus, 1.2 x10^7^ HEK-293T cells were plated onto a 15cm tissue culture dish. Twenty-four hours after plating, cells were co-transfected with the lentiviral transfer vector (EYFP^β5t^FOXN1^CMV^) and the psPAX2 and pMD2.G viral packaging vectors at a ratio of 4:3:2 using PEI-Pro (PolyPlus Transfection) following the manufacturer’s protocol. Culture media was exchanged 6 hours post-transfection with 10mL of fresh DMEM (Thermo Fisher Scientific) complemented with 10% FBS, non-essential amino acids (NEAA), penicillin/streptomycin and L-glutamine. Lentiviral supernatant was collected at 48- and 72-hours post transfection, pooled and filtered with a 0.45 µm cellulose acetate syringe filter (Sartorius), subsequently overlaid onto 5 mL of 20% sucrose and ultracentrifuged at 24,000 rpm for 2.5 hrs (Beckman XPN80, SW32.Ti). The viral pellet was resuspended in PBS.

For the generation of the FOXN1-EYFP reporter cell line, 4D6 cells were virally transduced with the EYFP^β5t^FOXN1^CMV^ expression vector. A range of multiplicity of infection (MOIs) was used to determine the virus titer required to achieve a 5% transduction efficiency to ensure a single integration of the construct in successfully transduced cells. Transduction was performed in the presence of 8µg/mL hexadimethrine bromide (Polybrene). The resulting stable cell line was evaluated 72 hours post transduction for EYFP expression levels using flow cytometry. Transduced cells were repetitively sorted by FACs to obtain a population with at least 75% EYFP positive cells as successfully transduced cells displayed the tendency to decrease the frequency of EYFP positive cells over time.

### Cloning of sgRNAs in the Cas9 expression vector and CRISPR mediated deletion of selected genes in the EYFP^β5t^FOXN1^CMV^ 4D6 reporter cell line

Single guide RNAs (sgRNAs) targeting FOXN1, YBX1 and CBP were selected from the GeneScript database https://www.genscript.com/gRNA-detail/1387/CREBBP-CRISPR-guide-RNA.html.

The following guide sequences were used

sgFOXN1: F:5’CACCGTGCTCGTCATTTGTGTCCGA3’

R:5’AAAC TCGGACACAAATGACGAGCA C3’,

sg CBP F:5’CACCGGAATCACATGACGCATTGTC3’

R: 5’AAACGACAATGCGTCATGTGATTCC3’

sg YBX1 F:5’CACCGGGACCATACCTGCGGAATCG3’

R:5’AAACCGATTCCGCAGGTATGGTCCC3’. Each pair of guides was cloned into the Cas9-2A-mRuby2 vector (https://www.addgene.org/110164/ Addgene #110164) following the protocol of the Feng Zhang laboratory (https://media.addgene.org/cms/filer_public/e6/5a/e65a9ef8-c8ac-4f88-98da-3b7d7960394c/zhang-lab-general-cloning-protocol.pdf). A single colony of transformed bacteria per guide was selected, picked, grown overnight in liquid cultures, miniprepped and eventually sent for sequencing to confirm the successful ligation into the Cas9-2A-mRuby2 vector.

The CRISPR mediated deletion of selected genes was performed by transfecting 4D6 EYFP^β5t^FOXN1^CMV^ cells with the respective guides using Fugene. Three technical replicates were prepared per sample. Cells transfected only with the EYFP^β5t^FOXN1^CMV^ vector, only with Fugene, or the sgCCR5 (a guide targeting a gene irrelevant for FOXN1 function) served as negative controls. Forty-eight hours post-transfection cells were collected in FACS buffer (1x PBS, 5% FCS), stained with DAPI for live/dead cell discrimination and analysed by flow cytometry. Live cells, DAPI negative, were gated for mRuby2 positivity (detected at 561/564nm) to select successfully transduced cells. Finally, the frequency of EYFP positivity was determined for cells that are mRuby2 positive.

### Mice

Animals were maintained under specific pathogen-free conditions and experiments were performed according to institutional and UK Home Office regulations. Mouse colonies were maintained at the University of Oxford Biomedical Science facilities. Age and gender matched wild-type mice were used in all experiments as a reference for genetically modified animals.

Mice heterozygous for a *Foxn1* allele with a single nucleotide loss at position 1470 (designated FOXN1^WT/Δ505^) were generated at the Genome Engineering Facility of the MRC Weatherall Institute of Molecular Medicine, University of Oxford, using the CRISPR/Cas9 assisted mESC targeting. In detail, murine JM8.1 C57B/6J ES cells were transfected with the pX459 plasmid construct that allowed expression of a Puromycin resistance gene, the Cas9 endonuclease and a single guide RNA (sgRNA) targeting the region in which the base pair change was expected to take place (Addgene 62988). The single stranded donor oligonucleotide (ssODN) for homology directed repair was also co-transfected to allow homologous integration of the desired genetic change. The ssODN was complementary to exon 8 of the mouse *Foxn1* locus with a single base pair deletion of an adenine at cDNA position 1370, and it also contained a *de novo* restriction enzyme site (PvuII) that was inserted in the sequence without changing the codon usage. Successfully targeted mouse embryonic stem cell (mESC) clones, verified by cloning and Sanger-sequencing, were injected into the blastocysts of albino C57BL/6BrdCrHsd-Tyrc mice. Resultant chimeric animals were selected by the presence of black coat color and confirmed by genotyping. Male chimeras were mated with albino C57BL/6BrdCrHsd-Tyrc female mice to establish germline transmission. Successful integration of the mutant allele was ultimately verified by genotyping. Heterozygous *Δ505* mice were mated with C57BL/6J WT black mice to expand the colony. For genotyping, the FOXN1^WT/Δ505^ mice, genomic DNA was extracted from ear clips and amplified by polymerase chain reaction (PCR) using the relevant forward (FP: 5’AAACTGGGCTCTCCGCTGCTG3’) and reverse (RP:5’AGTAGAGTATCGTGCATGGTCCTGG3’) primers (Taq polymerase (Sigma) at an annealing temperature (Tm) 64°C. PCR amplicons were digested using PvuII (NEB) restriction enzyme at 37°C for 1 hour and were run on an agarose gel. The undigested PCR amplicon (uncut) was detected at 389bp size. PvuII digested amplicons from FOXN1^WT/WT^ mice generated two bands of 389bp and 70bp size respectively, whereas PvuII digested amplicons from FOXN1^WT/Δ505^ heterozygous mice generated two additional bands of 213bp and 97bp sizes respectively

FTP^FOXN1^ mice: to probe Foxn1 promoter activity, C57BL/6 mice were rendered transgenic for the expression of a fluorescent timer protein (FTP) with a slow conversion kinetic [41] under the transcriptional control of the endogenous Foxn1 locus. In brief, the mCherry variant was inserted upstream of endogenous ATG start codon in exon 2. To engineer the targeting vector, homology arms were generated by PCR using BAC clone RP24-306F3 and RP23-204B15 from the C57BL/6J library as templates. In the targeting vector, the Neo cassette was flanked by self-deletion anchor sites. DTA was used for negative selection of mESC clones successfully targeted.

### Thymic epithelial cell and thymocyte isolation

Adipose and other tissues were removed from isolated thymi which were subsequently subjected to three rounds of enzymatic digestion for which cut up lobes were incubation at 37°C in Liberase digestion buffer (PBS Gibco 1X, Liberase Roche 2.5 mg/ml, DNase Roche 10 mg/ml). Cells were incubated with anti-CD45 beads (Miltenyi Biotec) as per manufacturer’s recommendations and subjected to CD45 depletion using the “depleteS” program on the AutoMACS separator (Miltenyi Biotec). CD45 depleted fractions were stained in PBS supplemented with 2% FBS (both Sigma) for flow cytometric analysis.

For thymocyte analysis, cells were obtained by gentle disruption of thymic lobes using frosted glass slides (Thermo Fisher Scientific). Cells were filtered through a 70µm filter (Greiner) and washed with ice-cold PBS supplemented with 2% FBS before being stained for downstream flow cytometric analysis using the reagents listed in the below table. Extracellular antibody stains were conducted for 20min at 4°C in the dark. CCR7 staining was performed at 37°C water-bath for 30min. The UEA-1 lectin (Vector Laboratories) was used pure, followed by anit-UEA1 biotin staining with secondary streptavidin-BV605.For the identification of dead cells, the Live/Dead Fixable Aqua stain kit (Thermo Fisher Scientific) was used. For flow cytometric phenotyping and cell sorting, a BD FACS Aria III (BD Biosciences) was used.

---

**Figure S1:**
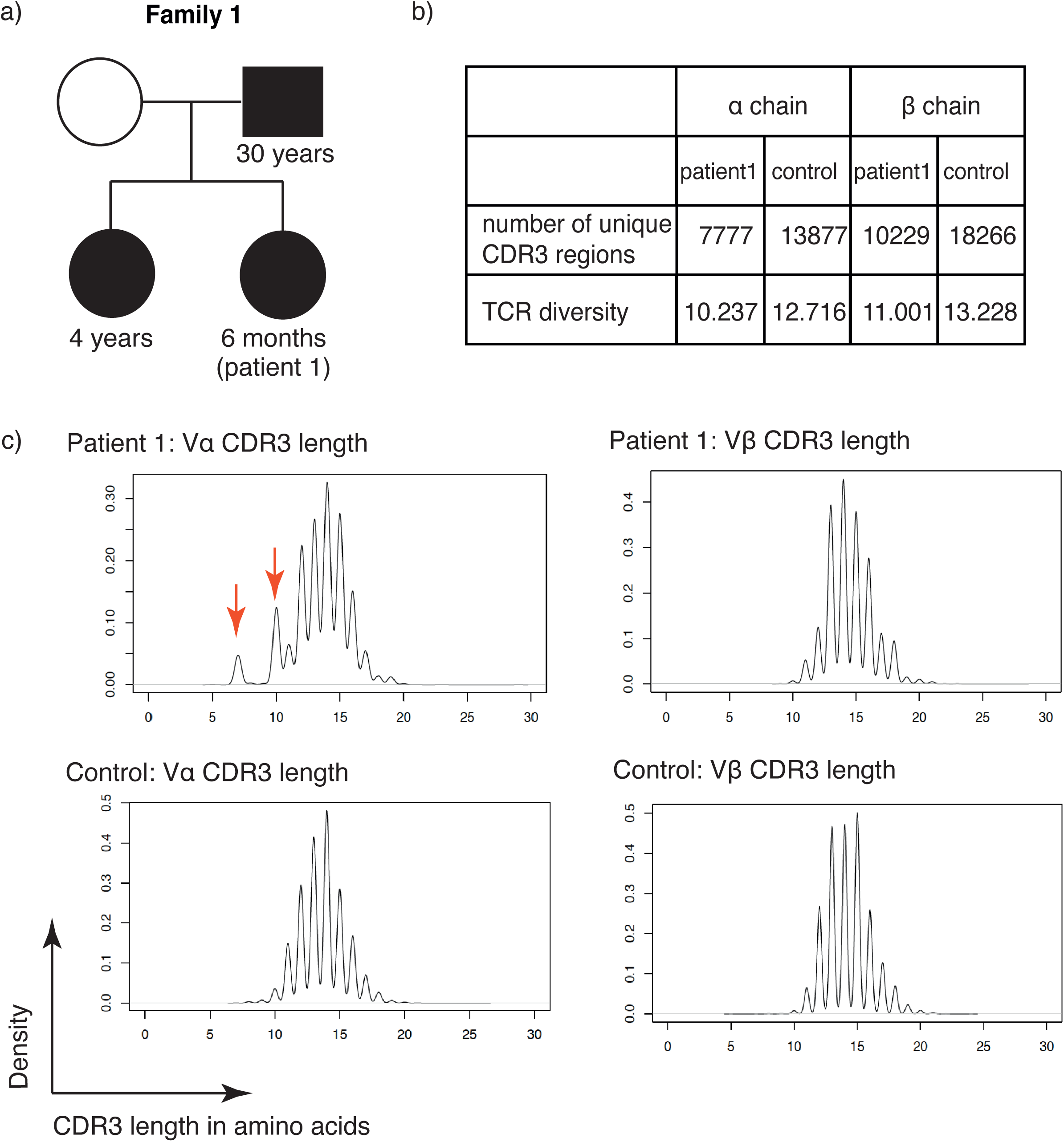
Clinical and experimental data of patients with novel FOXN1 variants. (a) Family pedigree of the index patient with the Δ550 FOXN1 mutation. Squares represent male and circles female family members. Filled symbols indicate individuals with the Δ550 FOXN1 mutation. Arrow identifies the index patient. Whole genome sequencing on isolated peripheral blood mononuclear cells from the index patient, father and older sister revealed a common single base pair deletion in exon 7 of FOXN1 (c.1370delA, p.H457FS*93; see also panel e). Other chromosomal rearrangements or abnormalities in the patients were excluded by Comparative Genomic Hybridization array (CGH). Moreover, other known genetic variants associated with primary immunodeficiencies (PID) were not found in any of the individuals. The index patient was initial brought to medical attention due to severe dermatitis, a history of chronic diarrhea and an acute chest infection caused by rhinovirus. Peripheral blood testing showed a T cell lymphopenia (32% of total lymphocytes; normal range 51–77%. Supplementary Table 1) with absent T cell receptor excision circles (TRECs). A chest ultrasound revealed the absence of a thymus. Immunological diagnostics demonstrated normal T cell proliferative responses to phytohemagglutinin as well as normal total serum IgG levels. The 4-year-old sister also had a reduction in naïve T cells (Supplementary Table 1), absent TRECs and a missing thymus but was otherwise thriving and clinically well. The 30-year-old father also lacked TRECs, but he was clinically well. Both the father and the sister also had high Epstein-Barr virus (EBV) serum loads but neither had developed any further symptoms. Following the initial clinical presentation, the index patient remained well and did not require further medical attention. It is of note that neither alopecia nor nail dystrophy were noted in the index patient, the older sister or their father. (b) Reduced alpha and beta chain TCR diversities compared to an age-matched control. The table shows the total number of unique CDR3 regions and TCR diversity as calculated by the Shannon entropy. Low Shannon indices reveal lower diversity [65](c) CDR3 length in amino acids of TCR alpha and beta chains of peripheral T cells isolated from patient 1 and an age-matched control as assessed by MiSeq. The red arrows indicate oligoclonal expansion of T cells with CDR3 length of 7 and 10 amino acids respectively.

**Figure S2:**
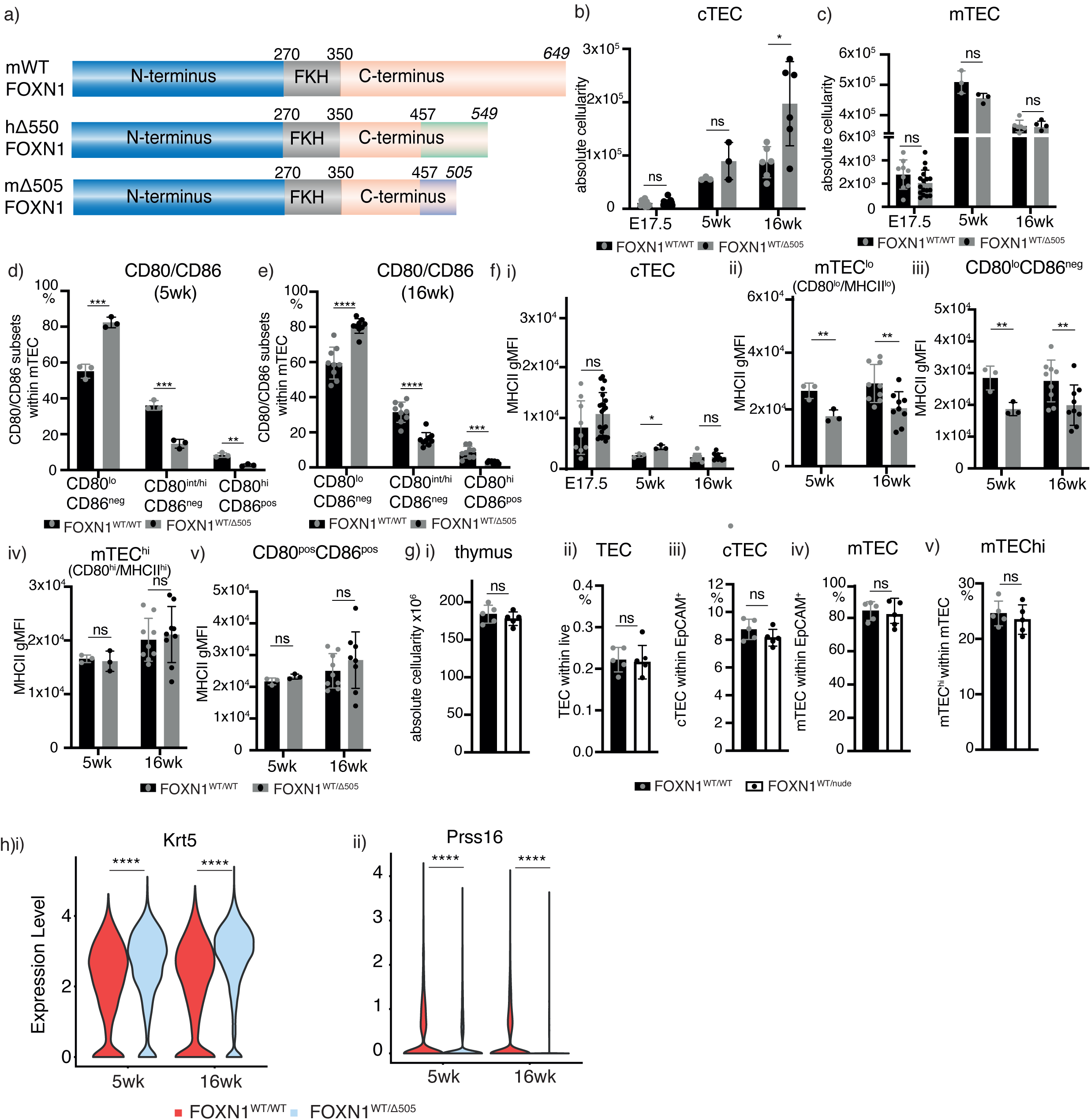
Mice heterozygous for the Δ505 mutation exhibit a partial block in TEC differentiation and differences in gene enrichment. Schematic representation of mouse wild-type (mWT), human Δ550 (Δ550) and mouse Δ505 (mΔ505) FOXN1. FKH: forkhead domain. Green and blue boxes represent the scrambled in the human and mouse FOXN1 sequence, respectively. Numbers in italics indicate the position of the stop codon. (b-e) Analysis of FOXN1^WT/WT^ and FOXN1^WT/Δ505^ mice at the indicated ages for cTEC, mTEC, and CD80^+^CD86^+^ mTEC cellularity. (f) Level of MHC cell surface expression on (i) cTEC, (ii) MHC^low^ mTEC, (iii) CD80^low^CD86^-^ mTEC, (iv) MHC^high^ mTEC and (v) CD80^+^CD86^+^ mTEC of FOXN1^WT/WT^ and FOXN1^WT/Δ505^ mice at the indicated ages. (g) Cellularity of 5 week old male FOXN1^WT/nude^ mice: (i) total thymus cellularity and frequencies of (ii) total TEC, (iii) cTEC, (iv) mTEC, and (v) mature (i.e. MHCH^high^) mTEC. (h) Violin plots with expression levels of (i) *Krt5*, (ii) *Prss16* at 5 and 16-week-old intertypical TEC. Data is from 4 (E17.5), 1 (week 5) and 2 (week 16) independent experiments with at least 3 FOXN1^WT/WT^ and n = 3 FOXN1^WT/Δ505^ male mice for (b-e and g). (f) Data is from 1 independent experiment with 5 FOXN1^WT/WT^ and 5 FOXN1^WT/nude^ male mice. (h) data is from 3 FOXN1^WT/WT^ and 3 FOXN1^WT/^Δ505 mice at 5 and 16 weeks of age, (h) 22,228 interypical TEC analysed. Mean and SD is indicated 0.05 (ns), 0.0332(*), 0.0021(**), 0.0002 (***), two-tailed unpaired t-test for (b-g), p<0.0001(****).

**Figure S3:**
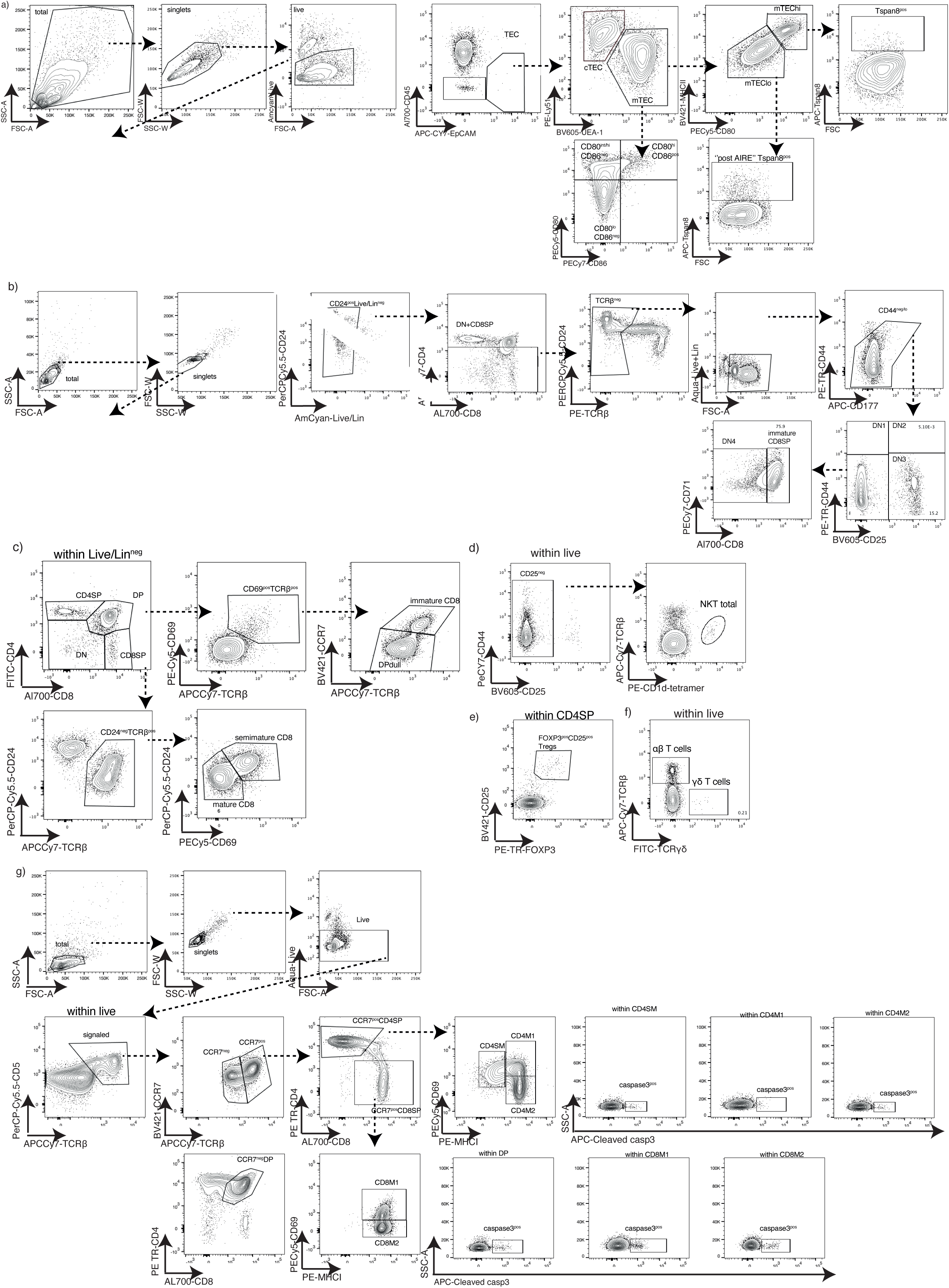
Gating strategy for the identification of TEC and thymocyte subpopulations by flow cytometry. (a) – (g) Representative flow cytometric plots of wild-type mice at 16 weeks of age to identify the indicated cell populations. Lineage-negativity was defined as the absence of cell surface expression of CD11b, CD11c, Gr1, CD19, CD49b, F4/80, NK1.1, TCRγδ, and Ter119.

**Figure S4:**
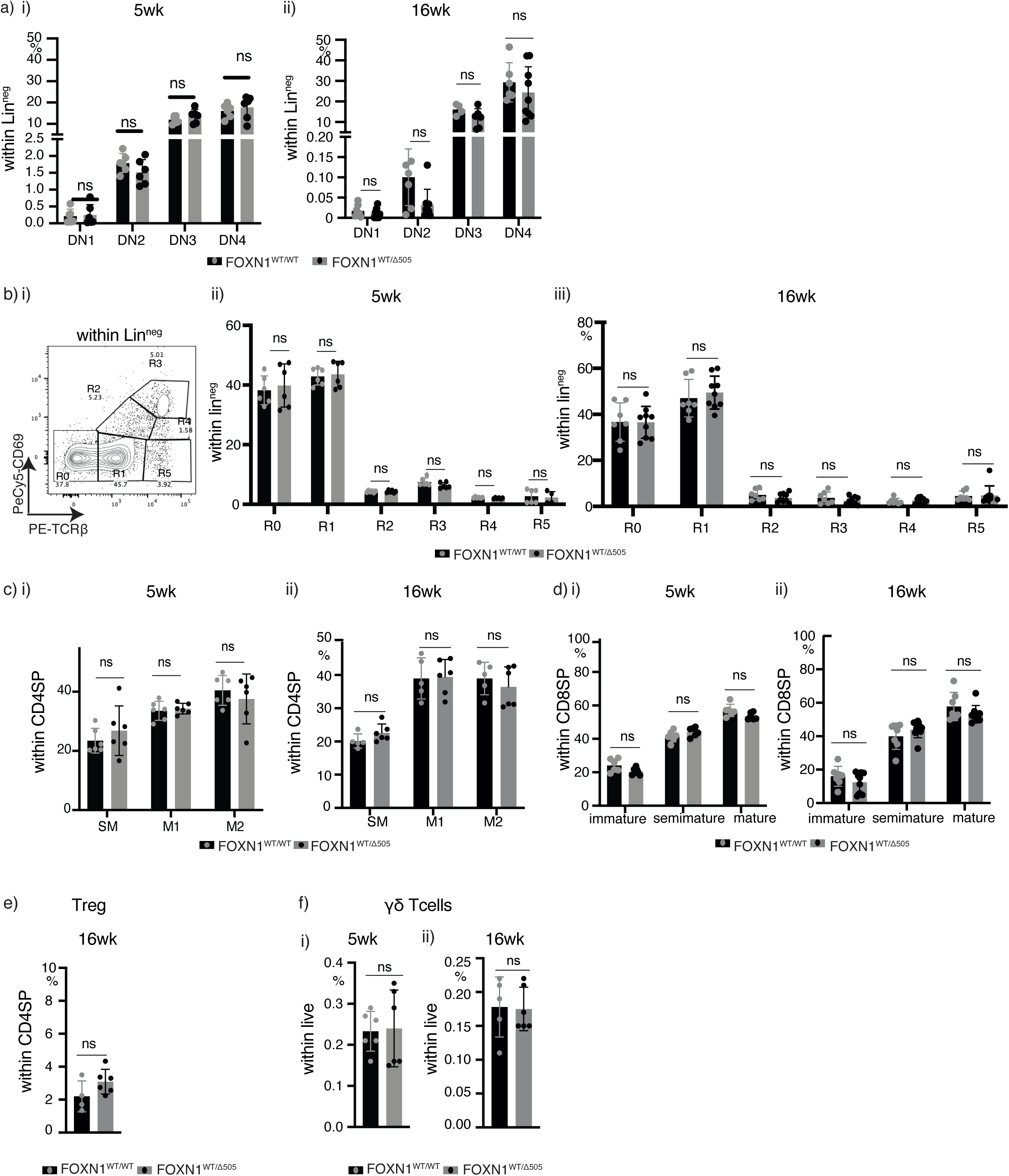
Thymocyte development in mice heterozygous for Δ505 FOXN1. Thymocyte analysis in 5 and 16 week old FOXN1^WT/WT^ and FOXN1^WT/Δ505^ mice. (a) Lineage-negative (Lin-) CD4^-^CD8^-^ (double negative, DN) thymocytes were stained for their cell surface expression of CD44 and CD25 defining 4 separate subpopulations: DN1 (CD44^+^CD25^-^), DN2 (CD44^+^CD25^+^), DN3 (CD44^-^CD25^-^) and DN4 (CD44^-^CD25^-^). Lin- was defined as negative for the expression of CD11b, CD11c, Gr1, CD19, CD49b, F4/80, NK1.1, TCRγδ, and Ter119. (b) Thymocyte maturational stages based on TCR and CD69 cell surface expression. (i) Dot plot analysis of wild-type thymocytes indicating the chosen gates R0-R5. Frequency of indicated thymocyte subpopulations. (c) Stages of CD4SP thymocytes. The CD4SP cells were grouped into semi-mature (SM: CD69^+^MHC I^low^), mature 1 (M1: CD69^+^MHC I^+^) and mature 2 (M2: CD69^-^ MHC I^+^) thymocytes [51]. (d) Stages of CD8SP thymocytes. The CD8SP cells were grouped into immature (CCR7^pos^, TCRβ^pos^, CD69^pos^ within DP), semimature (CD24^pos^, CD69^pos^ within CD8SP) and mature (CD24^neg^, CD69^neg^ within CD8SP) thymocytes. (e) Frequency of regulatory T cells (Treg: FOXP3^+^CD25^+^ CD4SP) (f) Frequency of γδ T cells. Each symbol represents data from an individual wild-type or mutant mouse at the indicated age. Data is from 3 (panels a,d) and 2 independent experiments (panels c, e, and f). Mean value and SD are shown and were calculated by two-tailed unpaired t-test. p-values: ≥0.05 (ns). The flow cytometric gating strategies are shown in Supplementary Figure 3.

**Figure S5:**
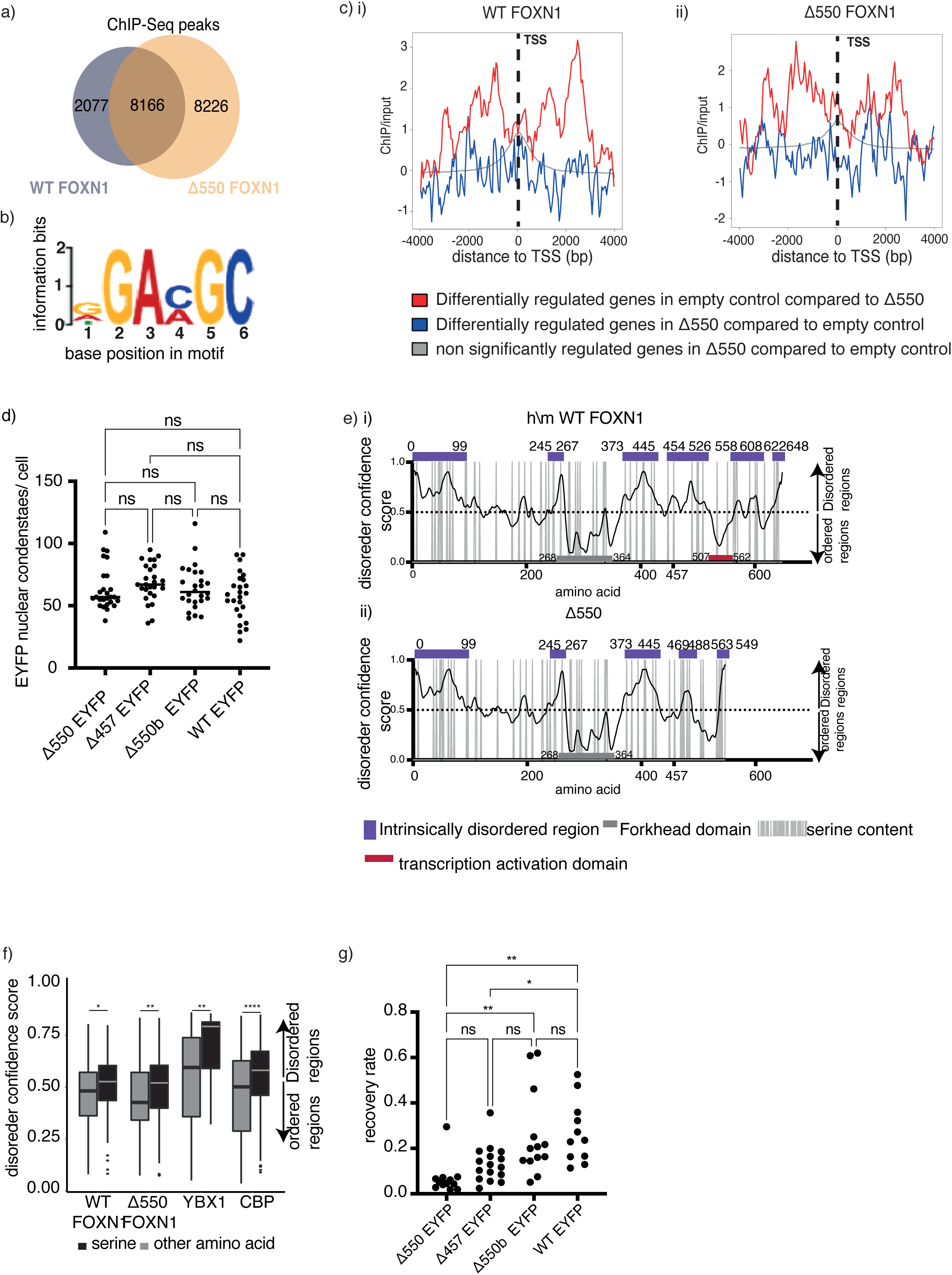
Molecular characteristics of FOXN1 variants. (a) ChIP-Seq peaks of wild-type and Δ550 FOXN1 expressed in 4D6 TEC. The Venn diagram shows the number of ChIP-Seq peaks that are identified by only wild-type, only Δ550 FOXN1 and both. (b) Weblogo of the MEME-derived FOXN1-binding site motif for TSS-associated peaks (–5 kb before and 100 bp after TSS), E = 1.7 x 10^-362^ for the WT peaks and E=2.5 x 10^-258^or the Δ550 FOXN1 peaks. (c) Enrichment of (i) wild-type FOXN1 and (ii) Δ550 FOXN1 ChIP-Seq signals over input in a 4kb region flanking the TSS for genes differentially regulated by wild-type FOXN1. All experiments in (a-c) were performed in biological triplicate (although, as detailed in the methods section, one of the Δ550 samples was excluded from the RNA-sequencing analysis as an outlier (d) quantification of nuclear condensates formed by EYFP-labelled Δ550, Δ457, Δ550b, wild-type FOXN1 variants. (e) Graphs showing intrinsic disorder regions and serine content for WT (i) and Δ550 FOXN1(ii). PrDOS (Protein DisOrder prediction System) disorder confidence score is shown on the y axis and amino acid positions on the x-axis. Confidence score above 0.5 predicts a disordered region. Vertical gray lines designate the presence of a serine at a given amino acid position. The purple bar designates the IDRs, the grey bar the forkhead domain and the red bar the transcriptional activation domain. (f) Box plot showing distribution of serine across ordered and disordered protein regions within WT, Δ550, CBP and YBX1 proteins. (g) Fluorescence’s recovery rate [arbitrary units (a.u.)/sec] of EYFP-labelled Δ550, Δ457, Δ550b, wild-type FOXN1 nuclear condensates following photobleaching. Mean and SD is indicated, (a-b) unpaired t-test 0.05 (ns), 0.0332(*), 0.0021(**).

**Figure S6:**
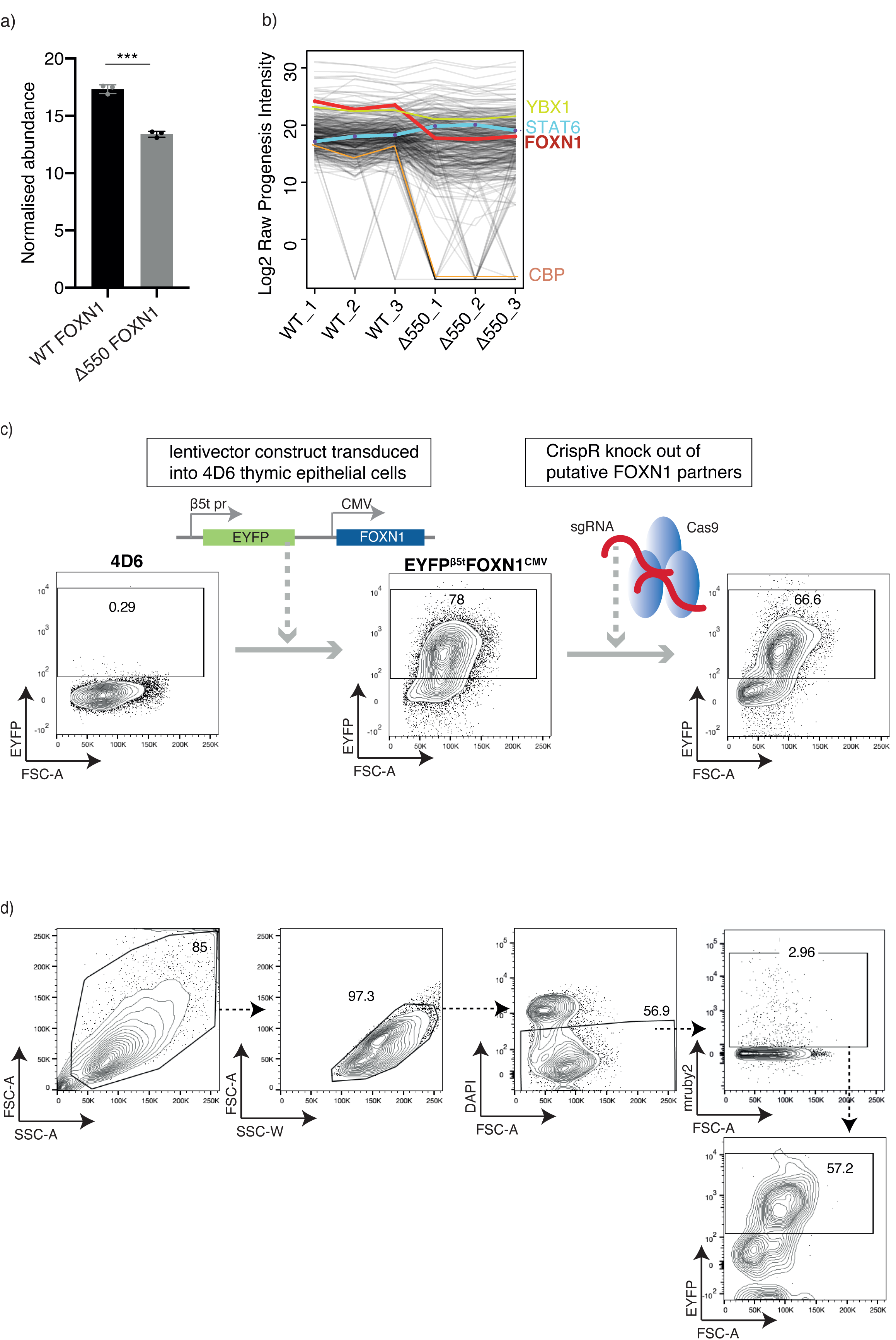
Characterisation of FOXN1 interacting partners. (a) Protein abundance of wild-type or Δ550 FOXN1 in anti-flag pull downs in lysates from 4D6 cells transfected to express wild-type (WT) or Δ550 FOXN1 tagged with a flag. The immunoprecipitates were analysed by Liquid Chromatography with tandem mass spectrometry (LC-MS-MS). Each point corresponds to an individual sample. The data is from a single experiment with three biological replicates per condition. Data shows the mean value and SD. (b) Normalised abundance of proteins identified by immunopreciptation and Liquid Chromatography with tandem mass spectrometry (LC-MS-MS). Comparison of 3 samples of 4D6 cells each transfected to express wild-type (WT) and Δ550 FOXN1 showing the normalized abundance of FOXN1 (red line), STAT6 (blue line), CBP (orange line), and YBX1 (green line). (c) Schematic representation of the generation of the 4D6-EYFP^β5t^ FOXN1^CMV^ reporter cell line used for the validation of candidate proteins interacting with wild-type FOXN1. 4D6 cells were stably transduced to constitutively express FOXN1 under the CMV promoter that in turn controls the transcription of EYFP under the transcriptional control of the *Psmb11* promoter which is a direct target of FOXN1 and drives β5t protein expression. (d) Flow cytometric gating strategy for the identification of EYFP positivity in live 4D6-EYFP^β5t^ FOXN1^CMV^ cells that have been transfected with guide RNAs as identified by mruby2 positivity.

**Figure S7:**
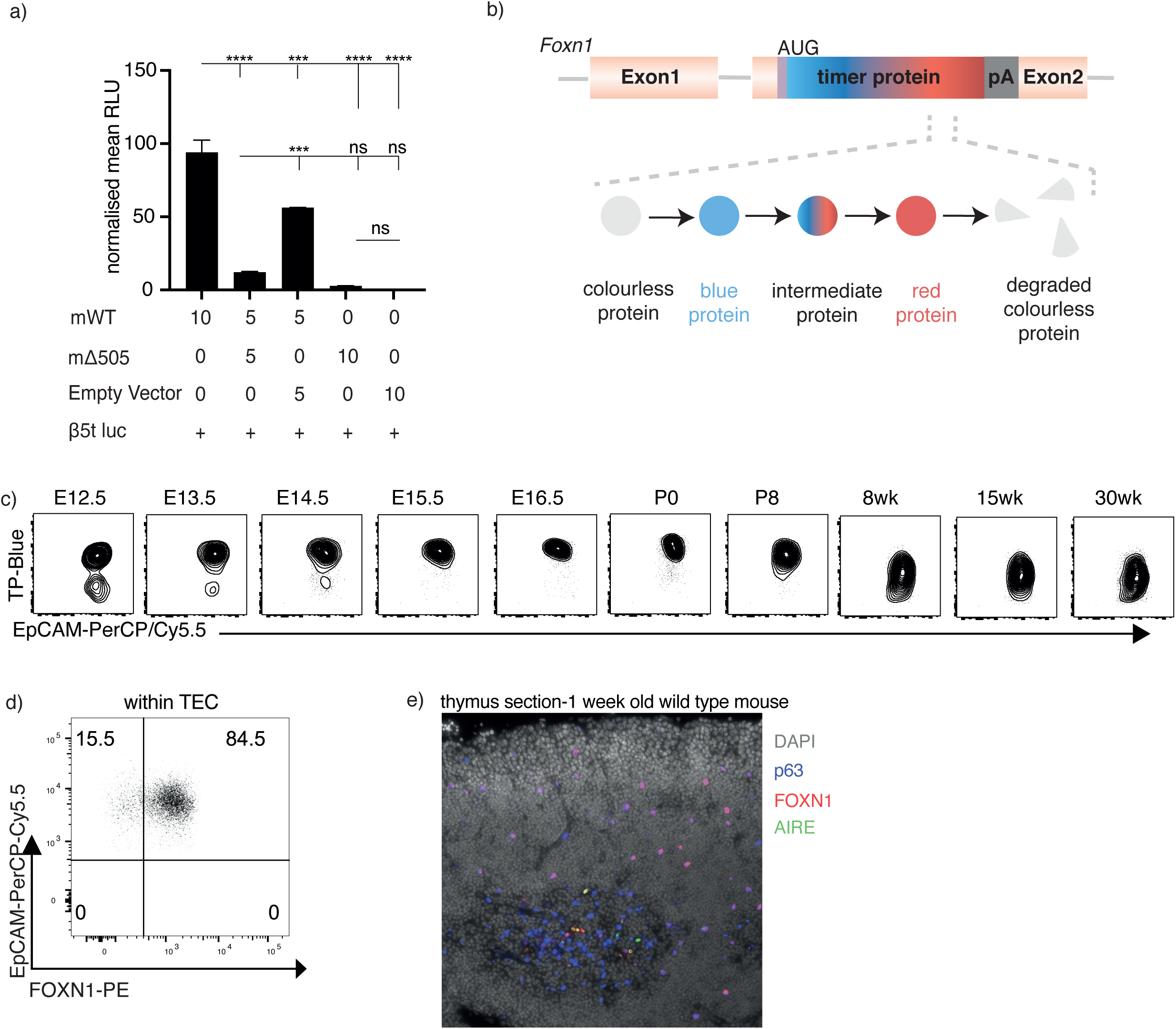
FOXN1 function and expression. (a) Expression of a luciferase reporter under the transcriptional control of the *Psmb11* promoter (β5t luc) in TEC1.2 cells transfected with indicated expression vectors. Constitutive renilla expression was used in each transfectant as an internal control and reporter activity was measured as relative light units (RLU) following correction. mWT: mouse wild-type, m Δ505: mouse Δ505 FOXN1. The data is from 3 independent experiments with each 3 technical replicates. Mean value and SD are shown and were and statistically compared by two-tailed unpaired t-test: p-values: ≥0.05 (ns), <0.0001(****). (b) Schematic representation of the timer protein construct. (c) representative flow cytometry plots of FTP-blue versus EpCAM in total TEC isolated from the FTP^FOXN1^ mice (d) representative flow cytometry plots of EpCAM versus FOXN1 in total TEC isolated from wild-type mice. Mean value and SD are shown in bar graphs and were calculated with two-tailed unpaired t-test; 0.05 (ns), 0.0332(*), 0.0021(**), 0.0002 (***).(e) Immunofluorescent analysis of thymic section from 1 week old wild-type mice for the expression of FOXN1, AIRE and p63. DAPI was used to counterstain cell nuclei.

**Table S1:** Gene ontology analysis of biological processes in the indicated TEC subtypes. Enrichment in FOXN1^WT/WT^ TEC. Data is from 3 FOXN1^WT/WT^ and 3 FOXN1^WT/^Δ505 mice at 5 and 16 weeks of age, with 22,228 interypical TEC analysed.

**Table S2:** Gene ontology analysis of biological processes in the indicated TEC subtypes. Enrichment in in FOXN1^WT/Δ505^ TEC. Data is from 3 FOXN1^WT/WT^ and 3 FOXN1^WT/^Δ505 mice at 5 and 16 weeks of age, with 22,228 interypical TEC analysed.

**Table S3:** List of putative FOXN1 binding partners. Uniprot KB was used to determine the subcellular localisation and the gene ontology (GO) of each protein. VRK3 and STAT6 are the two proteins that inversely correlate with the amount of FOXN1.

**Table S4:** list of antibodies used for the FACs phenotypic analysis and FACs cell sorting.

