## Supplementary material for "A FOXN1 mutation competitively displaces wild-type FOXN1 from higher-order nuclear condensates to cause immunodeficiency": Table S3

| Accession | Gene Name | Subcellular location [CC] | Gene ontology |
|---|---|---|---|
| P17096 | HMGA1 | Nucleous | DNA Binding [GO:0003677], negative regulation of transcription [GO:0045892]; positive regulation of transcription [GO:0045944]; |
| P12956 | XRCC6 | Nucleous | DNA Binding [GO:0003677]; positive regulation of transcription [GO:0045944]; |
| Q92793 | CBP | Nucleous | acetyltransferase activity [GO:0016407]; RNA polymerase II transcription factor binding [GO:0001085]; transcription coactivator activity [GO:0003713]; transcription corepressor activity [GO:0003714]; transcription factor binding [GO:0008134]; DNA Binding [GO:0003677] |
| P06748 | NPM | Nucleous | activating transcription factor binding [GO:0033613]; RNA binding [GO:0003723]; transcription coactivator activity [GO:0003713]; positive regulation of transcription by RNA polymerase II [GO:0045944]; |
| Q14103 | HNRPD | Nucleous, cytoplasm | mRNA splicing GO:0000398, |
| Q13283 | G3BP1 | Nucleous, cytoplasm | DNA Binding [GO:0003677] |
| P78527 | PRKDC | Nucleous | DNA Binding [GO:0003677], positive regulation of transcription [GO:0045944]; |
| Q7KZF4 | SND1 | Nucleous | RNA binding [GO:0003723]; transcription coregulator activity [GO:0003712] |
| Q8NC51 | PAIRB | Nucleous | RNA binding [GO:0003723] |
| P68104 | EF1A1 | Nucleous | translation [GO:0006412]; |
| P67809 | YBX1 | Nucleous, cytoplasm | mRNA splicing GO:0000398, negative regulation of transcription [GO:0000122]; transcription by RNA polymerase II [GO:0006366] |
| Q6AHZ1 | Z518A | Nucleous | DNA Binding [GO:0003677] |
| Q12906 | ILF3 | Nucleous | DNA binding [GO:0003677]; |
| P04908 | H2A1B | Nucleous | DNA binding [GO:0003677]; |
| O15353 | FOXN1 | Nucleous | DNA Binding [GO:0003677] |
| Q96AE4 | FUBP1 | Nucleous | positive regulation of gene expression [GO:0010628]; transcription by RNA polymerase II [GO:0006366], DNA Binding [GO:0003677] |
| P09651 | ROA1 | Nucleous | mRNA splicing GO:0000398 |
| Q05639 | EF1A2 | Nucleous | translation [GO:0006412]; |
| Q96T23 | RSF1 | Nucleous | negative regulation of transcription [GO:0045892]; positive regulation of transcription by RNA polymerase II [GO:0045944]; acetyltransferase activity [GO:0016407]; |
| P43243 | MATR3 | Nucleous | RNA binding [GO:0003723] |
| P04792 | HSPB1 | Nucleous | RNA binding [GO:0003723] |
| O75150 | RNF40 | Nucleous | mRNA 3'-UTR binding [GO:0003730]; |
| Q9NZI8 | IF2B1 | Nucleous | RNA binding [GO:0003723] |
| P11940 | PABP1 | Nucleous | mRNA splicing GO:0000398 |
| P52272 | HNRPM | Nucleous | mRNA splicing GO:0000398 |
| P51991 | ROA3 | Nucleous | mRNA splicing GO:0000398 |
| Q8N684 | CPSF7 | Nucleous | mRNA splicing GO:0000398 |
| P48634 | PRC2A | Nucleous | RNA binding [GO:0003723] |
| P31942 | HNRH3 | Nucleous | mRNA splicing GO:0000398 |
| Q9H869 | YYAP1 | Nucleous | regulation of cell cycle [GO:0051726] |
| Q8IV63 | VRK3 | Nucleous | ATP binding [GO:0005524]; protein phosphatase binding [GO:0019903]; protein serine/threonine kinase activity [GO:0004674] |
| P42226 | STAT6 | Nucleous | DNA-binding transcription activator activity [GO:0001228]; [positive regulation of transcription [GO:0045944]; |
