## Supplementary material for "A FOXN1 mutation competitively displaces wild-type FOXN1 from higher-order nuclear condensates to cause immunodeficiency": Table S4

| Antibody | Clone | Conjugate | Source |
|---|---|---|---|
| CCR7 | 4B12 | BV421 | BD Biosciences |
| CD11b | M1/70 | biotin | BioLegend |
| CD11c | N418 | biotin | BioLegend |
| CD177 (c-kit) | 2B8 | APC | Biolegend |
| CD19 | 1D3 | biotin | eBioscience |
| CD1d | tetramer | PE | University of Birmingham |
| CD24 | M1/69 | APC | BioLegend |
| CD24 | M1/69 | PerCP eFluor710 | eBioscience |
| CD25 | PC61.5 | BV605 | BioLegend |
| CD25 | PC61.5 | BV421 | eBioscience |
| CD4 | RM4-5 | PE Texas Red | BioLegend |
| CD4 | RM4-5 | APC-Cy7 | BioLegend |
| CD4 | RM4-5 | FITC | BioLegend |
| CD44 | IM7 | PE Cy7 | eBioscience |
| CD44 | IM7 | PE Texas Red | eBioscience |
| CD45 | 30-F11 | Alexa Fluor 700 | BioLegend |
| CD49b | DX5 | biotin | BioLegend |
| CD5 | 53-7.3 | PerCP, Cy5.5 | BioLegend |
| CD69 | H1.2F3 | PE Cy5 | BioLegend |
| CD71 | RI7217 | PE Cy7 | BioLegend |
| CD80 | 16-10A1 | PE Cy5 | BioLegend |
| CD86 | GL-1 | PE Cy7 | BioLegend |
| CD8a | 53-6.7 | Alexa Fluor 700 | BioLegend |
| Cleaved casp 3 (Asp175) | D3E9 | AF647 | Cell signaling |
| EPCAM | G8.8 | APC-CY7 | BioLegend |
| EPCAM, | G8.8 | PerCP Cy5.5 | BioLegend |
| F4/80 | BM8 | biotin | BioLegend |
| FOXN1 | - | PE | a kind gift by HR Rodewahl[24] |
| FoxP3, | FJK-16s | eF450 | eBioscience |
| Gp2 | 2F11-C3 | FITC | MBL |
| Gr1 | RB6-8C5 | biotin | BioLegend |
| H2Kb (MHCI) | AF6-88.5 | PE | BioLegend |
| Ly51 | 6C3 | PE | BioLegend |
| MHC-II | 28-14-8 | BV421 | eBioscience |
| NK1.1 | PK136 | biotin | BioLegend |
| Streptavidin | - | BV605 | BioLegend |
| TCRβ | clone H57-597 | APC-Cy7 | BioLegend |
| TCRβ | H57-597 | PE | eBioscience |
| TCRβ | H57-597 | FITC | eBioscience |
| TCRγδ | UC7-13D5 | FITC | BioLegend |
| TCRγδ | GL3 | biotin | eBioscience |
| TER119 | TER119 | biotin | BioLegend |
| Tspan8 | 657909 | APC | R&D Systems |
